# Charting the brain networks of impulsivity: Meta-analytic synthesis, functional connectivity modelling and neurotransmitter associations

**DOI:** 10.1101/2023.07.04.547631

**Authors:** Martin Gell, Robert Langner, Vincent Küppers, Edna C. Cieslik, Theodore D. Satterthwaite, Simon B. Eickhoff, Veronika I. Müller

## Abstract

Impulsivity is a multidimensional construct that plays a crucial role in human behaviour and is believed to be a transdiagnostic marker of several psychiatric disorders. However, given its multifaceted nature, comprehensive investigations of its neural correlates are challenging. In this study, we used a multi-modal approach to investigate the functional network organisation of two domains in which impulsivity manifests: decision-making and action control. Within domain ALE meta-analyses identified two distinct functional systems: one located in the default-mode network, associated with value-based judgments and goal-directed decision-making, and the other distributed across higher-order networks associated with cognitive control. These systems were organised into four specialised communities of default-mode, cingulo-insular, frontoparietal, and temporal regions. Integration within those communities was associated with serotonin receptor density. These findings reinforce insights from previous behavioural research and provide substantial evidence for the multidimensional nature of impulsivity on the neural level.

## Introduction

Impulsivity has been defined as the “predisposition toward rapid, unplanned reactions to internal or external stimuli without regard to the negative consequences of these reactions to the individual or to others” (Moeller, Barratt, Dougherty, Schmitz, & Swann, 2001). Impulsive behaviours are a pervasive part of life for many individuals, from reckless driving (Teese & Bradley, 2008) or reactive aggression (Gvion & Apter, 2011) to smoking (Sharma, Markon, & Clark, 2014) or thrill-seeking (Whiteside & Lynam, 2001; Cyders et al., 2007). Thus, impulsivity plays a crucial role in the human condition, being strongly intertwined with cognitive control and decision-making (Dalley, Everitt, & Robbins, 2011). Heightened impulsivity is believed to be a hallmark of several psychiatric disorders such as attention-deficit/hyperactivity disorder (ADHD), substance abuse, and bipolar disorder (Moeller et al., 2001), which has informed theories of impulsivity as a transdiagnostic marker (Berlin & Hollander, 2014; Amlung et al., 2019). Therefore, understanding the neural mechanisms behind impulsivity is of high research and societal value.

Behavioural and theoretical investigations of impulsivity indicate it is a multidimensional psychological construct (Bari & Robbins, 2013; Caswell, Bond, Duka, & Morgan, 2015; Dalley et al., 2011; Dalley & Robbins, 2017; Dick et al., 2010; MacKillop et al., 2016; Reynolds, Ortengren, Richards, & de Wit, 2006; Reynolds, Penfold, & Patak, 2008), and some authors even argue there is no single umbrella construct of impulsivity at all (Cyders, 2015; Strickland & Johnson, 2020). Proposed models vary, resulting in a lack of consensus on the number and characteristics of the constituent dimensions. Furthermore, trait-based models of impulsivity such as the UPPS-P (Cyders et al., 2007; Whiteside & Lynam, 2001) often appear to be unrelated to assessments of behavioural performance, yielding largely independent bodies of evidence (Sharma et al., 2014; Strickland & Johnson, 2020). Within most performance-based models, impulsivity is believed to manifest as suboptimal decision-making due to discounting of delayed consequences and the failure to inhibit prepotent response tendencies. The subjective decrease in reward value as a function of the delay in obtaining that reward has been labelled as delay consequence sensitivity (DCS; (Strickland & Johnson, 2020), ‘impulsive choice’ (Hamilton, Mitchell, et al., 2015; Winstanley, Eagle, & Robbins, 2006) or ‘impulsive decision-making’ (Sharma et al., 2014). It is classically investigated using the delay discounting paradigm. Failures of response inhibition, the capacity to inhibit a prepotent response tendency, have been otherwise referred to as ‘impulsive action’ (Winstanley et al., 2006) or ‘rapid-response impulsivity’ (Hamilton, Littlefield, et al., 2015). In humans, it is typically investigated using go/nogo, stop-signal or 5-choice serial reaction time tasks.

In the last few decades, there has been broad interest in understanding the neural mechanisms of impulsivity, with studies reporting associations with brain activity (Christakou, Brammer, & Rubia, 2011; Sripada, Gonzalez, Phan, & Liberzon, 2011; DeVito et al., 2013; Wilbertz et al., 2014; Wang, Shen, et al., 2017; Anandakumar et al., 2018), connectivity (Wang, Zhou, et al., 2017; H. Cai, Chen, Liu, Zhu, & Yu, 2020) and neurochemistry (de Boer & Koolhaas, 2005; Winstanley, Theobald, Dalley, & Robbins, 2005; Koffarnus, Newman, Grundt, Rice, & Woods, 2011). However, given the multidimensional nature of impulsivity, comprehensive investigations of its neural correlates are challenging. Moreover, results across studies are difficult to compare, as the primary outcome measures are often correlations with impulsive traits that show little overlap with performance-based assessments of impulsivity. As impulsive individuals display altered behavioural responses on DCS and response inhibition tasks, neural correlates can be investigated already on the level of activity in those behavioural tasks without the need for correlational analyses. Here, many studies have explored task-based brain activations during response inhibition (Aron & Poldrack, 2006; Sebastian et al., 2013) and DCS (McClure, Laibson, Loewenstein, & Cohen, 2004) as well as associations with DCS-related processes such as the subjective valuation of rewards (Kable & Glimcher, 2007). Partially echoing behavioural findings, these studies point to two largely distinct functional systems associated with response inhibition and DCS (Bari & Robbins, 2013; Dalley et al., 2011), with potentially overlapping regions across domains within the prefrontal cortex (PFC; (Schüller, Kuhn, Jessen, & Hu, 2019; Noda et al., 2020). Specifically, evidence suggests that the multiple-demand network, sometimes referred to as the frontoparietal or cognitive control network, subserves inhibitory control exerted to prevent premature responding (Duncan, 2010; Cieslik, Mueller, Eickhoff, Langner, & Eickhoff, 2015; Zhang, Geng, & Lee, 2017). Conversely, the default-mode network (DMN) or the valuation system together with the dorsolateral prefrontal cortex is believed to underlie DCS (Owens et al., 2017; Schüller et al., 2019; Noda et al., 2020). However, direct comparisons of brain activation during response inhibition and DCS are lacking (see Wang et al. (2016) for a comparison with respect to resting-state connectivity and grey matter volume). Thus, the architecture of the functional brain networks linked to the different impulsivity dimensions and their interactions remains poorly understood.

A large body of theoretical work considers impulsivity as a form of a trade-off between self-control and impulsive systems (Whiteside & Lynam, 2001; Dalley et al., 2011; Sharma et al., 2014). Behaviourally, impulsive responding during tasks probing inhibition and discounting is characterised by commission errors or fast reaction time (Bari & Robbins, 2013; Ioannidis, Hook, Wickham, Grant, & Chamberlain, 2019) and steeper discounting of future rewards (Frost & McNaughton, 2017), respectively. However, given the trade-off between control and error-sensitive systems on the neural level in cognitive control (Duckworth, Potticary, & Badyaev, 2018), a comprehensive account of the neural mechanisms behind impulsivity ought to capture both regions related to ‘impulsive’ error responses as well as those linked to ‘controlled’ correct responses within each domain. In the literature, successful inhibition has been associated with the anterior insula, medial frontal cortex and right frontoparietal regions (Zhang et al., 2017; Cieslik, Ullsperger, Gell, Eickhoff, & Langner, 2023). Conversely, posterior medial frontal cortex covering pre-supplementary motor area (pre-SMA) and anterior midcingulate cortex (aMCC), dorsal posterior cingulate cortex, and thalamus are believed to play an important role in error monitoring as activity in these regions has been reliably found during errors of commission (Ullsperger, Fischer, Nigbur, & Endrass, 2014; Cieslik et al., 2023). Functional dissociations between impulsive and controlled responding during the delay discounting task are less clear. The ventral striatum, ventromedial PFC and anterior cingulate cortex (ACC) have been implicated in choices of smaller sooner (SS) rewards or steeper discounting; conversely, dorsolateral PFC and right parietal regions have been associated with choices of larger later rewards (LL) and shallower discounting (Schüller et al., 2019; Noda et al., 2020).

While neuroimaging evidence points to two largely distinct functional systems associated with response inhibition and DCS, investigations on the neurochemical level are more mixed, with several neurotransmitter systems believed to play a significant role in modulating impulsivity (Chamberlain & Sahakian, 2007; Dalley et al., 2011; Dalley & Robbins, 2017). Psychostimulant drugs used to treat ADHD such as methylphenidate block the reuptake of dopamine and norepinephrine. In most patients, they substantially reduce symptoms and can improve response inhibition even in healthy individuals (Aron & Poldrack, 2006; Hanwella, Senanayake, & de Silva, 2011; Nagashima et al., 2014). Functionally, these improvements may be partly ascribed to increased right inferior frontal and insula activation (Rubia et al., 2014). In rodents, atomoxetine, a selective norepinephrine reuptake inhibitor reduces delay discounting and enhances inhibition (Robinson et al., 2008). Outside psychostimulants, dopamine is classically associated with addiction (Berke & Hyman, 2000; Wise & Robble, 2020) and has been suggested as a major candidate for passing reward prediction errors within the valuation system (Nasser, Calu, Schoenbaum, & Sharpe, 2017). Findings from the animal literature show that lesions to the nucleus accumbens - a dopamine-rich nucleus - increase impulsivity on DCS tasks and may also impair response inhibition (Basar et al., 2010). Finally, there is some evidence for the involvement of serotonin in response inhibition, which is impaired following serotonin depletion (Worbe, Savulich, Voon, Fernandez-Egea, & Robbins, 2014). It has also been inversely related to aggression, a behavioural manifestation of impulsivity, with serotonin 5HT1A/1B receptor agonists reducing aggressive behaviour(de Boer & Koolhaas, 2005; Duke, Bègue, Bell, & Eisenlohr-Moul, 2013; da Cunha-Bang & Knudsen, 2021).

Here we aimed to delineate a comprehensive brain network associated with impulsivity using coordinate-based ALE meta-analyses (Turkeltaub, Eden, Jones, & Zeffiro, 2002; Eickhoff et al., 2009; Eickhoff, Bzdok, Laird, Kurth, & Fox, 2012) to synthesise the pertinent neuroimaging literature. We focus on two cognitive-behavioural dimensions that show consensus across most performance-based models of impulsivity and are most commonly investigated in the neuroimaging literature: delayed consequence sensitivity and response inhibition. To this end, we investigate both the activity associated with impulsive responding (commission errors or choices of SS rewards) and non-impulsive, ‘controlled’ responding (successful inhibition or choices of LL rewards) within each dimension. To capture other relevant processes involved in the execution of the delay discounting task, we also included associations with subjective value (Kable & Glimcher, 2007). Next, we characterised the network organisation using resting-state functional connectivity and graph-theoretical methods in two independent large-scale datasets to uncover the functional architecture of the impulsivity networks. Finally, given the widespread use of neurotransmitter-acting medication to treat conditions with impulsive symptoms (Chamberlain & Sahakian, 2007), we investigated if the impulsivity network function could theoretically be modulated by neurochemistry. To this end, we explored associations between network organisation and receptor density of neurotransmitter systems associated with impulsivity (dopamine, serotonin and norepinephrine) obtained from PET imaging.

## Results

### Meta-analysis

#### Delayed-Consequence Sensitivity

Analysis of experiments investigating DCS revealed significant findings only for impulsive responding (i.e. impulsive decision-making: choices of SS over LL and correlation with discounting factor k) and subjective value (correlation and parametric modulation) contrasts. Impulsive responding (Fig. 1A) led to consistent activation of the ventromedial prefrontal cortex (VMPFC), left frontal pole (FP), ACC and bilateral ventral caudate extending to the nucleus accumbens hereafter referred to as ventral striatum (VS) (Haber, 2011). Analysis of experiments correlating activity with subjective value revealed convergence in a largely overlapping network (Fig. 1B). Conjunction analysis revealed that left VMPFC, bilateral VS and right ACC were common in both meta-analyses. Conversely, contrast analyses showed that only FP was specific to impulsive responding, while subcallosal cingulate cortex (scACC) and posterior cingulate cortex (PCC) were specific to subjective value (see Supplementary Figures 1 and 2). There were no converging clusters for experiments testing controlled responding (choices of LL over SS). The exclusion of studies that correlated measures of impulsivity such as the discount rate k (thus including only the ‘pure’ SS > LL and LL > SS contrasts) revealed similar results (see Supplementary Figure 3).

**Figure 1.**
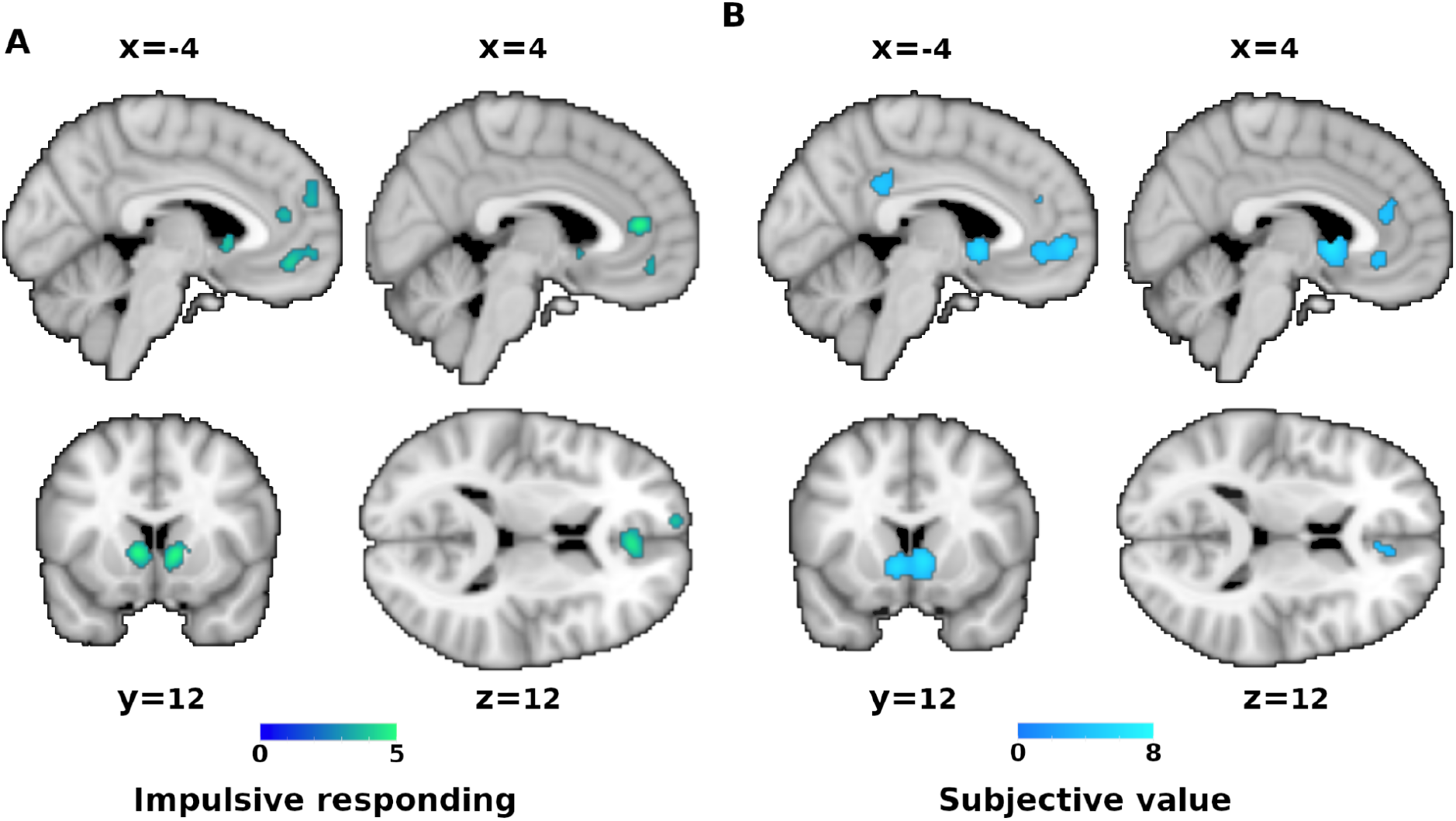
Delayed-consequence sensitivity. Results of the meta-analysis on brain activity correlates of (A) impulsive responding and (B) subjective value. Colour codes z-score.

#### Response Inhibition

Due to the low number of published experiments that directly contrasted impulsive and controlled responding (i.e. inhibition failure vs. success), a meta-analysis of this contrast was not possible (Eickhoff et al., 2016). We, therefore, computed two meta-analyses of experiments examining brain activation during failed or successful inhibition against baseline and subsequently tested for differences as well as commonalities between them on the meta-analytic level (see Methods for further details). Results of the individual analyses of failed or successful inhibition against baseline can be found in supplementary figure 4. These analyses revealed a widespread network of insular, frontoparietal and subcortical regions in line with previous findings (Cieslik et al., 2023).

The meta-analytic contrast analysis revealed stronger convergence for impulsive responding (failed vs. successful inhibition) in preSMA, aMCC, the right anterior section of the superior frontal gyrus (aSFG) and right supramarginal gyrus (SMG) (Fig. 2A orange-yellow). Stronger convergence for controlled responding (successful vs. failed inhibition) was found across the lateral frontal and dorsal premotor cortex (dPMC) in addition to the right temporal and parietal regions, right anterior insula (aI) and left putamen (Fig. 2A blue). Figure 2B illustrates the conjunction analysis across the meta-analyses of failed and successful inhibition against baseline.

**Figure 2.**
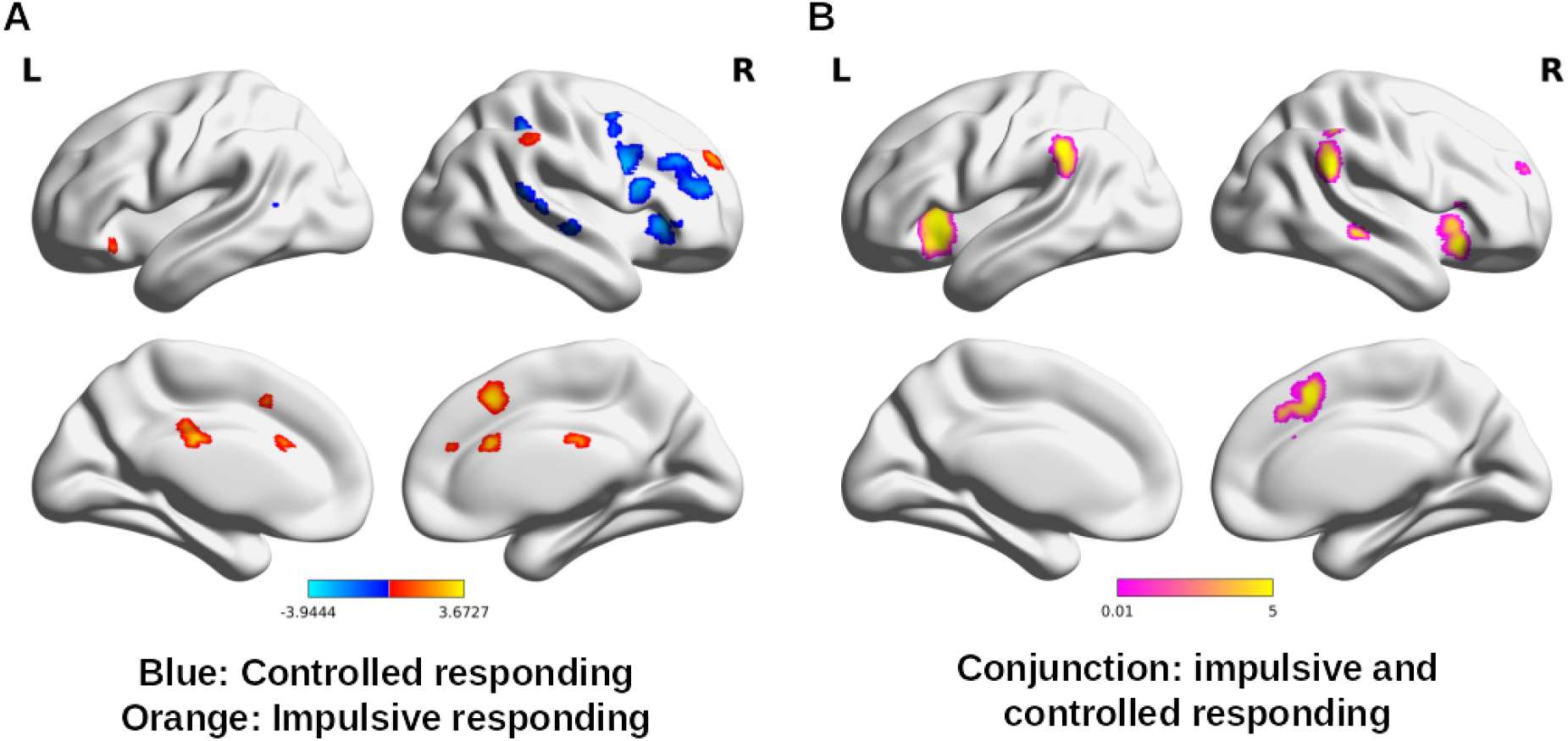
Response Inhibition. Results of (A) meta-analytic contrast and (B) conjunction analyses of successful inhibition > baseline and failure of inhibition > baseline contrast meta-analyses.

#### Network characterisation

To explore the functional organisation of the resulting meta-analytic regions, we investigated their functional connectivity profiles, community structure and neurochemical properties. The nodes that were used for these analyses are displayed in Figure 3. For MNI coordinates and complete regions labels see Table 1. An overlay with Yeo et al.’s (2011) resting state networks (Fig. 3B) shows that nodes from the DCS meta-analyses were primarily located within medial DMN. Combined controlled and impulsive responding nodes were mostly found in the dorsal attention network, while the remaining nodes were distributed over frontoparietal and ventral attention networks (Fig. 3B). The extracted meta-analytic nodes and all result maps are available in the ANIMA database: https://anima.fz-juelich.de/studies (Reid et al., 2016).

**Figure 3.**
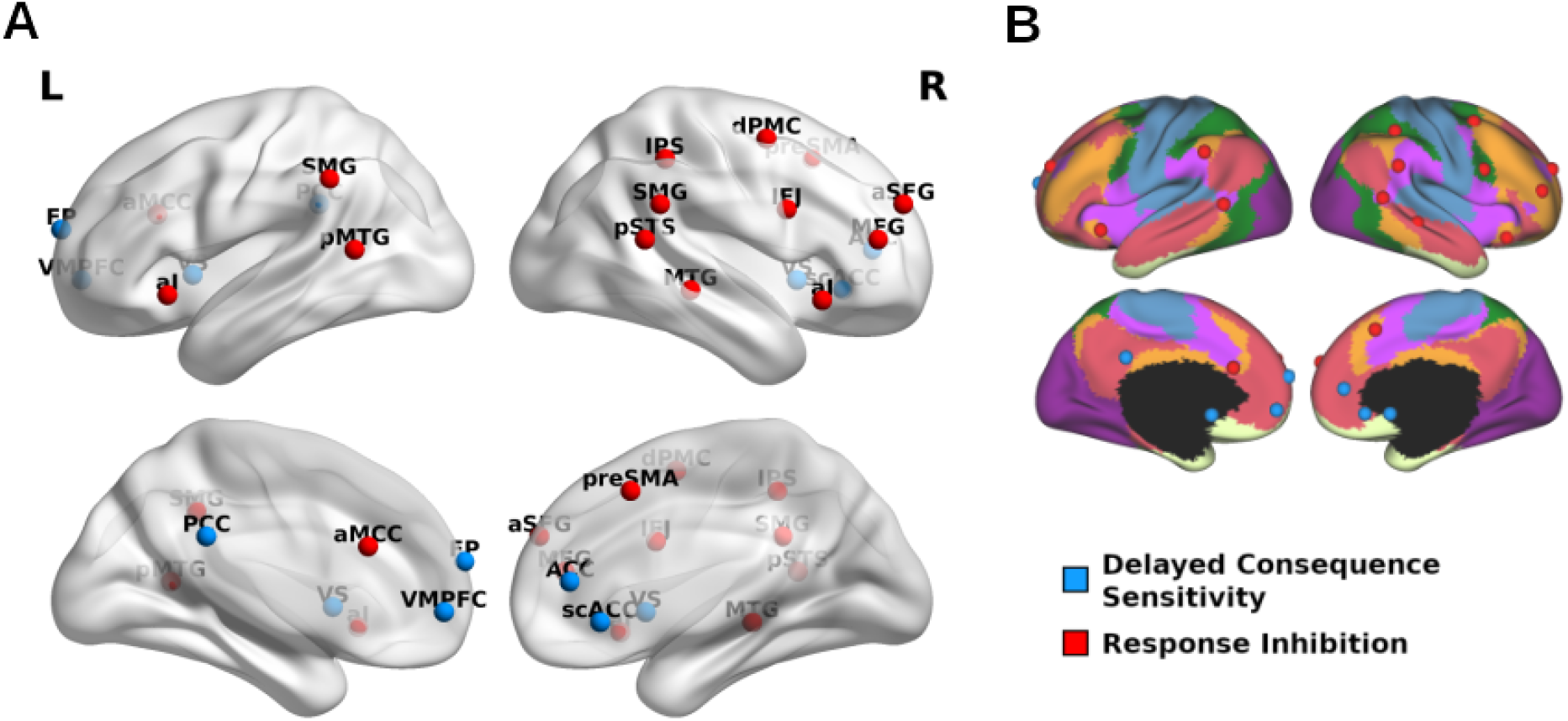
Nodes of the Impulsivity Network. (A) impulsivity network nodes: Delayed consequence sensitivity in blue and response inhibition in red. Panel (B) displays Impulsivity network nodes overlaid over Yeo et al., (2011) resting-state networks: visual (purple), somatomotor (blue), dorsal attention (green), ventral attention (pink), limbic (white), frontoparietal (orange) and default mode (red) networks.

**Table 1.** Meta-analytic nodes

| Region (abbreviation) | Hemisphere | MNI Coordinate |  |  |
| --- | --- | --- | --- | --- |
|  |  | x | y | z |
| Ventral striatum (VS) | Left | -8 | 10 | 0 |
| Ventromedial prefrontal cortex (VMPFC) | Left | -4 | 44 | -8 |
| Ventral striatum (VS) | Right | 10 | 12 | -2 |
| Anterior cingulate cortex (ACC) | Right | 8 | 42 | 10 |
| Frontal pole (FP) | Left | -10 | 62 | 18 |
| Subcallosal cingulate cortex (scACC) |  | 2 | 30 | -6 |
| Posterior cingulate cortex (PCC) |  | -2 | -38 | 28 |
| Anterior insula (AI) | Left | -38 | 20 | -8 |
| Anterior insula (AI) | Right | 32 | 22 | -10 |
| Pre-supplementary motor area (preSMA) | Right | 4 | 18 | 48 |
| Supramarginal gyrus (SMG) | Right | 60 | -42 | 28 |
| Supramarginal gyrus (SMG) | Left | -60 | -44 | 36 |
| Middle temporal gyrus (MTG) | Right | 54 | -30 | -6 |
| Superior frontal gyrus (SFG) | Right | 24 | 54 | 28 |
| Anterior midcingulate cortex (aMCC) |  | 0 | 24 | 24 |
| Inferior frontal junction (IFJ) | Right | 48 | 8 | 26 |
| Middle frontal gyrus (MFG) | Right | 40 | 44 | 14 |
| Posterior superior temporal sulcus (pSTS) | Right | 58 | -48 | 14 |
| Intraparietal sulcus (IPS) | Right | 40 | -40 | 46 |
| Posterior middle temporal gyrus (pMTG) | Left | -58 | -52 | 12 |
| Dorsal premotor cortex (dPMC) | Right | 38 | 0 | 54 |

Abbreviations: MNI – Montreal neurological institute

#### Community Structure

To detect communities within the impulsivity network we used the Louvain community detection algorithm (Blondel, Guillaume, Lambiotte, & Lefebvre, 2008), which divides a network into non-overlapping groups of nodes. Using estimates of resting-state FC between all network nodes from 528 participants of the publicly available Nathan Kline Institute dataset (eNKI) (Nooner et al., 2012) as edges, this approach yielded a four-community solution (Fig. 4A). Repeating this procedure 1000 times, we observed a strong convergence across solutions suggesting that our four-community solution was not restricted to a local maximum in the solution space (Fig. 4B). To evaluate the robustness of our findings further, we repeated the community detection analysis using a different set of 316 unrelated subjects from the Human Connectome Project dataset (HCP) (Van Essen et al., 2013) and found an identical community structure (see Supplementary Figure 5).

**Figure 4.**
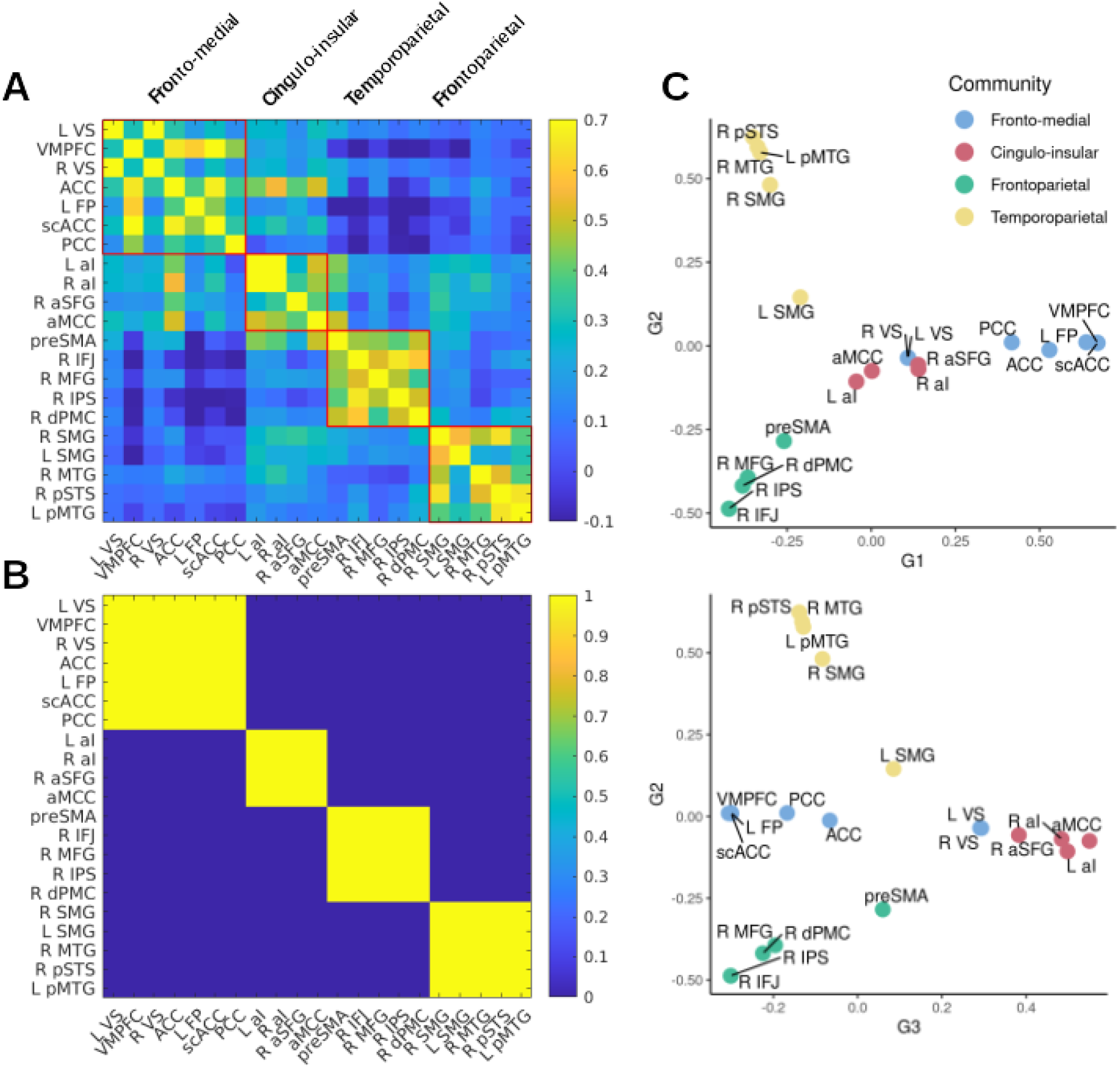
Impulsivity Network Communities. Panel A shows connectivity-based communities in the discovery sample (eNKI). The agreement matrix in panel B displays the consensus across 1000 repetitions of the community detection. Legend refers to the proportion of overlapping community solutions. Seed-voxel connectivity gradients are displayed in panel C. For a 3D depiction of the three components in C see: https://github.com/MartinGell/Impulsivity_networks

The first, fronto-medial community consisted of all DCS nodes in the network (VS, VMPFC, ACC, frontal pole, PCC and scACC). Regions related to response inhibition were subdivided into three different communities. In the order of appearance in Figure 4, the first of these comprised mostly regions of the so-called salience network (Seeley et al., 2007), i.e. bilateral aI and aMCC as well as right SFG. The next community spanned mainly right-lateralized frontoparietal regions (IFJ, MFG, dPMC and IPS) as well as preSMA. The last community consisted of temporoparietal regions (bilateral SMG, MTG, and pSTS). Interestingly, the cingulo-insular community was the only community to display positive coupling with regions of both the DCS and response inhibition networks.

Finally, we investigated the robustness of the resulting communities by a complementary whole-brain analysis. Here, principal component analysis of the pairwise similarity between maps of seed-to-voxel connectivity of the meta-analytic nodes was used to explore the dimensions along which they were organised in relation to the rest of the brain (for a scree plot, see Supplementary Figure 6). The initial three components that explained the most variance showed loadings that were in strong agreement with our community detection results suggesting the node-to-brain interactions paralleled node-to-node relationships (Fig. 4C). The first principal component showed that DCS nodes (except VS) displayed affinity in their connectivity with the rest of the brain while being dissimilar to the response inhibition regions. Similar properties were observed for the cingulo-insular, frontoparietal and temporoparietal communities along the second and third gradient revealing the closeness of within-community nodes in their whole-brain connectivity profiles. Results did not differ with varying sparsity or decomposition parameters.

#### Network Organisation Related to Receptor Density

Finally, we examined if network organisation was associated with neurotransmitters related to impulsivity across domains (Dalley & Robbins, 2017). In particular, given our systems approach, we were interested if the interactions between network nodes within and between communities are related to dopamine and serotonin receptor density as well as norepinephrine transporter density derived from PET imaging. Network organisation was assessed using two graph-theoretical measures: (i) within-module degree z-score, a measure of how well a node is connected to other nodes in its community and (ii) participation coefficient, a measure of how well a node is connected to other modules (Guimerà & Nunes Amaral, 2005). Only serotonin 5HT1a receptor density showed a positive relation to within-module degree z-score in both samples (eNKI: ρ = 0.49, p = 0.014; HCP: ρ = 0.64, p = 0.002), suggesting that node-wise serotonin expression was related to within-module integration (Fig. 5A).

**Figure 5.**
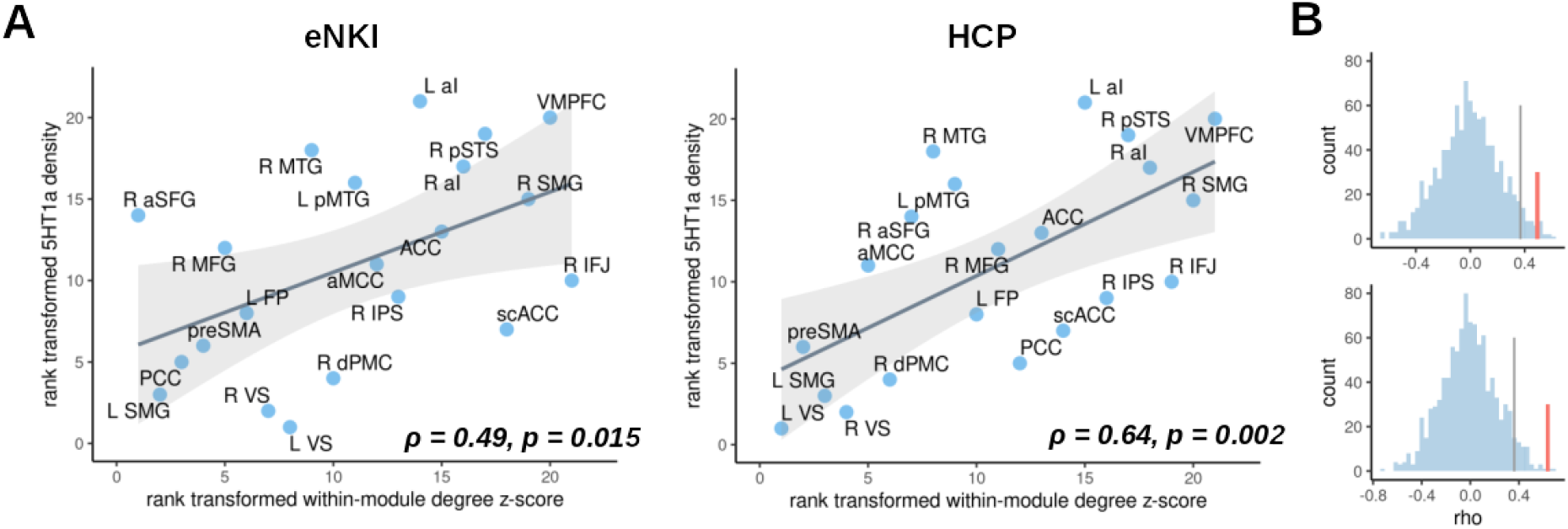
Association of community organisation with serotonin receptor 5HT1a. Panel A shows the relationship between within-module degree z-score (high scores indicate within-network integrator regions) and 5HT1a receptor in the discovery (left) and replication sample (right). Panel B displays permutation-derived null-distributions of correlation coefficients (Spearman’s rho) between receptor density and within-module degree z-score in the discovery sample (top) and replication sample (bottom). Observed correlation is marked with a red line and the significance level of 0.05 is indicated by a grey line.

**Figure 6.**
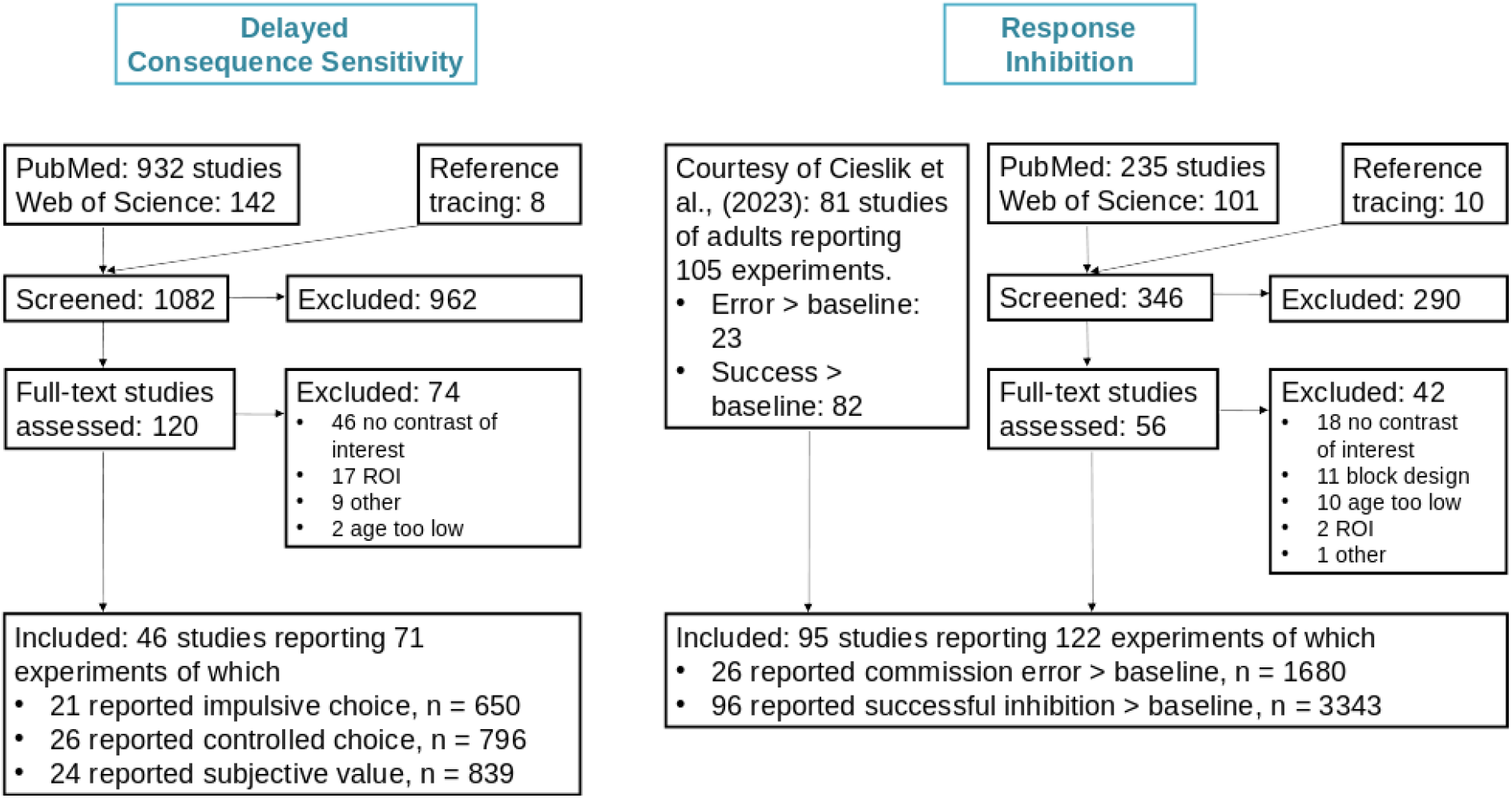
Flow Diagram of Study Selection for the Meta-analyses. The total sample size for each contrast is denoted with n. In the flow diagram, ‘study’ refers to a publication and by ‘experiment’ the specific contrast reported. In case a study reported multiple contrasts within the same category (e.g. stop signa), it was counted as one experiment.

## Discussion

The present study investigated brain networks associated with two dimensions of impulsivity: response inhibition and DCS. Using ALE meta-analyses of task-based fMRI studies we provide evidence for two distinct functional systems: one centred in the medial prefrontal cortex, ventral striatum and posterior cingulate cortex involved in DCS, and the second covering right lateral frontal cortex, temporoparietal regions, anterior insula and anterior midcingulate cortex subserving response inhibition. Community detection based on resting-state functional connectivity between all meta-analytically derived nodes in two large independent samples revealed four functional communities. The fronto-medial community included all DCS regions corroborating their dissociation from the other system. Response inhibition, in turn, was fractionated into three networks spanning frontoparietal, temporoparietal and cingulo-insular regions. Lastly, the integration of individual nodes within those communities calculated in two independent datasets was associated with serotonin receptor density.

### Two Systems

The results of our meta-analyses indicate that response inhibition and DCS domains of impulsivity differ not only in terms of their behavioural manifestations (Sharma et al., 2014; Stahl et al., 2014; MacKillop et al., 2016) but also on the neural level. Mirroring the behavioural dichotomy, we found two distinct sets of regions involved in each of the domains. The network of regions associated with DCS was mainly localised within the DMN (Raichle, 2015; Owens et al., 2017), while response inhibition covered the multiple-demand network (Duncan, 2010; Müller, Langner, Cieslik, Rottschy, & Eickhoff, 2015; Langner, Leiberg, Hoffstaedter, & Eickhoff, 2018). These findings directly support theoretical accounts proposing separate functional systems for individual impulsivity domains (Strickland & Johnson, 2020) and are in agreement with findings on delay discounting (Frost & McNaughton, 2017; Noda et al., 2020) and response inhibition (Zhang et al., 2017; Cieslik et al., 2023). The current work now provides a more fine-grained overview by differentiating controlled and impulsive processing within each dimension and considering both in the final network definition.

Within the framework of response inhibition, impulsivity has been described as an impairment in executive functioning (with inhibitory control being one of the major executive functions), while from the delayed-gratification perspective, impulsivity has been more associated with motivational processes that underlie decision-making (Bari & Robbins, 2013; Stahl et al., 2014). In line with this, the regions related to DCS identified here, especially vmPFC and ventral striatum, have been previously implicated in value-based decision-making (Haber & Knutson, 2010; Rangel & Hare, 2010). Similarly, regions related to response inhibition have been classically associated with executive functions (Duncan, 2010; Langner et al., 2018; Camilleri et al., 2018). These functional differences echo behavioural findings. Performance on response inhibition and delay discounting tasks are differentially related to treatment outcomes of impulsivity-related disorders (Sheffer et al., 2012; Stevens et al., 2014), impulsive behaviours such as reactive aggression or drug-taking (Sharma et al., 2014) and to pharmacological intervention (Winstanley, Dalley, Theobald, & Robbins, 2004; Worbe et al., 2014). For instance, after reviewing the literature, Stevens and colleagues (2014) concluded that retention and treatment success in addiction, a condition believed to be strongly related to impulsivity (de Wit, 2009; Dick et al., 2010), was likely related to performance in monetary incentive delay tasks, but not to commission errors in response inhibition. The present findings, therefore, show that behavioural differentiation between the two dimensions is also mirrored on the neural level by the involvement of two distinct neurocognitive systems.

### Four Communities

Activity within the response inhibition and DCS networks has been linked to both behavioural (Aron & Poldrack, 2006; Hariri et al., 2006; Wang et al., 2016) and clinical (Stevens et al., 2014) variability. However, to develop markers of psychopathology, interactions within and between large-scale systems are essential (Castellanos, Di Martino, Craddock, Mehta, & Milham, 2013; Bassett, Xia, & Satterthwaite, 2018). Moreover, co-occurring deficits in both response inhibition and steeper delay discounting within the same individual in conditions like addiction and ADHD are not uncommon (Bickel, Jarmolowicz, Mueller, Koffarnus, & Gatchalian, 2012; Castellanos-Ryan et al., 2014; Ioannidis et al., 2019; Yücel et al., 2019).

Thus, the two networks identified here cannot account for most impulsivity-related variability in isolation. A systems perspective that considers both within- and between-system interactions may be necessary to bridge this gap. To this end, we used resting-state functional connectivity between the meta-analytic nodes as well as between the nodes and the rest of the brain to identify their community organisation based on their intrinsic coupling patterns. Supporting our meta-analytic findings, the fronto-medial community comprised all DCS regions, suggesting tight integration. Conversely, response inhibition regions split into three communities (cingulo-insular, temporoparietal and frontoparietal) that strongly resemble previous reports (Camilleri et al., 2018; Langner et al., 2018).

The frontoparietal community corresponded to regions within the dorsal attention (IFJ, dPMC, IPS) and frontoparietal (preSMA, MFG) resting state networks (Yeo et al., 2011). The dorsal attention network is believed to subserve top-down control of visuospatial attention (Corbetta & Shulman, 2002) including attentional shifting (Kelley, Serences, Giesbrecht, & Yantis, 2008), while the preSMA has been implicated in cognitive control (Cole et al., 2013) and motor preparation (Kennerley, Sakai, & Rushworth, 2004). Directing attention to expected and relevant stimuli and intentionally enhancing the processing of these stimuli when they occur subserved by the DAN may thus enable the appropriate initiation or inhibition of actions when appropriate (such as when a stop or no-go signal appears). With the exception of the right MTG (located in the DMN), the temporoparietal community (bilateral SMG and STS) covered regions located in the posterior ventral attention network. The TPJ, which covers most of the community, has been argued to underlie contextual updating more generally (Geng & Vossel, 2013) and updating responses from action execution to action inhibition during the stop-signal task more specifically (Cieslik et al., 2015). Thus, inefficient updating or transfer of updated information to motor regions via preSMA may result in slower responses or failures of inhibition commonly observed in high-impulsive individuals (Bari & Robbins, 2013).

The last community displayed tight interactions between the anterior insula, aMCC and aSFG, which have been previously described as the salience network (SN) (Seeley et al., 2007; Gordon et al., 2017). The SN has been associated with detecting important or salient stimuli (Seeley et al., 2007), and is believed to initiate control signals and facilitate switching between higher-order networks (Menon & Uddin, 2010; Goulden et al., 2014). We observed positive associations between the cingulo-insular community and both the frontoparietal and temporoparietal communities supporting its role as a control element within the response inhibition network (for a similar account, see Camillieri et al., 2017). In action inhibition specifically, such top-down signals likely originate from the aMCC which has been previously linked to error monitoring (Ullsperger et al., 2014) and may be crucial to inhibitory planning in the preSMA that displayed a strong association with it. Taken together, by facilitating attention, control, updating and action planning, the three communities together likely produce the required behaviour: to enact or inhibit an impulsive response tendency.

The cingulo-insular community also displayed a positive association with the DCS subsystem. These results are in line with models of the salience network as a control element mediating the dynamic interactions between DMN and frontoparietal networks to facilitate goal-directed behaviour (Menon, 2011). Similarly, the cingulo-insular community may play a role in coordinating the fronto-medial and frontoparietal communities. Aberrant interactions between the frontoparietal networks, DMN and SN (i.e. the triple network model) (Menon, 2011) have been proposed to underlie a number of psychiatric disorders. It is thus not unlikely that impulsivity, itself a transdiagnostic marker (Berlin & Hollander, 2014), is related to the functional integrity of the cingulo-insular, fronto-medial and frontoparietal communities. Supporting this, connectivity between these large-scale systems has been already associated with discounting rate (Chen, Guo, Suo, & Feng, 2018), ADHD (W. Cai, Chen, Szegletes, Supekar, & Menon, 2018), addiction (Wang, Shen, et al., 2017; Zhang & Volkow, 2019) and impulsive symptoms in Parkinson’s disease (Koh et al., 2020). Similarly, findings of aberrant connectivity between the dlPFC (part of the frontoparietal subsystem) and ventral striatum (part of the delay sensitivity subsystem) in substance use disorder (Jollans et al., 2016; Ersche et al., 2020) and pathological gambling (Koehler et al., 2013) may be in part explained by a dysfunctional salience control subsystem. As such, inappropriate disengagement of either the frontoparietal or fronto-medial communities during task execution may result in apparent connectivity changes between them and affect behaviour (Liang, Zou, He, & Yang, 2016; Shine & Poldrack, 2018). Taken together, we propose the multidimensional construct of impulsivity is associated with a broad network including default-mode, frontoparietal, temporal and subcortical regions that can be distinguished into four communities. Interactions between these communities suggest that the entire network is ultimately involved in the final behavioural phenotype of impulsivity.

### Neurochemistry

To investigate the biological relevance of the identified community organisation, we explored the relationship between integration and segregation of the impulsivity network with the receptor/transporter density of three impulsivity-related transmitter systems of the brain. These analyses revealed that within-community integrator regions display a higher density of the serotonin 5HT1a receptor, suggesting that integration within communities may be modulated by available serotonin. Evidence of serotonin involvement in different impulsivity dimensions is mixed, with the strongest evidence implicating it in response inhibition (Dalley & Robbins, 2017). There is ample evidence for an inverse association between serotonin levels and aggression, a behavioural manifestation of impulsivity (Duke et al., 2013; Carhart-Harris & Nutt, 2017; da Cunha-Bang & Knudsen, 2021). Specifically, 5HT1A/1B receptor agonists have been shown to reduce aggressive behaviour in many species including humans (Cleare & Bond, 2000; Sperry, Thompson, & Wingfield, 2003; de Boer & Koolhaas, 2005; Popova, Naumenko, & Plyusnina, 2007), while a reduction in firing has been associated with increased aggression (Audero et al., 2013). Activation within regions that exhibited high within-community integration like the anterior insula and medial PFC has been previously proposed to regulate aggression (Blair et al., 2021). The present findings, therefore, indicate that serotonergic modulation of behaviours such as aggression might be associated with facilitated integration within communities. Interestingly, neither the norepinephrine transporter nor dopamine receptor density was found to be related to functional network organisation. Our results thus indicate that the mechanism of action of norepinephrine and dopamine on function may not be through altering network integration or segregation, warranting further investigation.

### Limitations and Outlook

The present investigation focused on neural responding during the execution of cognitive tasks measuring impulsivity. It, therefore, does not warrant any conclusions on the relationship between brain activity and self-report measures of impulsivity, as questionnaire-derived trait assessments often demonstrate limited correlations with performance-based assessments of impulsivity (Sharma et al., 2014). Future work may investigate whether the network identified here is affected by individual differences in trait impulsivity. Here we focused on the two best-characterised dimensions of impulsivity that were also most commonly investigated with fMRI. Some models suggest sustained attention (the ability to keep one’s attention focused over time) and risk-taking as additional components of impulsivity (Strickland & Johnson, 2020); however, there is substantial variance in proposed behavioural assessments. A meta-analysis of fMRI studies investigating sustained attention by Langner & Eickhoff (2013) has reported activations in regions largely overlapping with those identified here in the response inhibition network. Risky behaviours rarely play a substantial role in theoretical models of impulsivity and have been measured using the probability discounting task and Balloon Analog Risk Task (Lejuez et al., 2002). fMRI investigations during these tasks have revealed regions within the DCS network and parts of the multiple demand network (Peters & Büchel, 2009; Schonberg et al., 2012; Miedl, Peters, & Büchel, 2012; Seaman et al., 2018), suggesting overlapping activation with regions found in our meta-analyses. Therefore, the network described here may provide a largely comprehensive description of the the neurocircuitry associated with multidimensional construct of impulsivity.

### Conclusions

Taken together, our findings reinforce insights from previous behavioural research and provide substantial evidence for the multidimensional nature of impulsivity on the neural level. In particular, we identified and characterised two non-overlapping neurocognitive systems linked to processes underlying impulsive and controlled decision-making and action control. Each of these was centred in a distinct large-scale network of brain organisation. The first was located in the default-mode network associated with value-based judgements and goal-directed decision-making, the second was distributed across higher-order networks related to executive functions of action selection, planning and updating. These systems were found to be organised into four specialised communities of medial frontal, cingulo-insular, frontoparietal and temporal regions. Interactions between the communities and their coordination may affect the impulsivity of our behaviour and decision-making, with the modulation of community integration by serotonin emerging as a possible mechanism. Overall, our findings indicate that a transdiagnostic treatment of impulsivity (Berlin & Hollander, 2014) requires a marriage between symptoms, cognitive processes and functional systems that needs to be addressed in future research in order to advance our understanding of impulsivity and finally influence clinical practice. The research domain criteria framework of the NIH (Insel et al., 2010) has already taken steps in such a direction, with reward valuation and response selection/inhibition forming two separate components – but only the latter refers to impulsivity. Such developments, however, have yet to penetrate clinical research and practice.

## Methods

### Meta-analysis

#### Study selection

We performed a literature search using PubMed (https://www.ncbi.nlm.nih.gov/pubmed) and Web of Science (https://webofknowledge.com) for articles published until the 10th March 2021 that investigated brain activation related to either a DCS or response inhibition with fMRI or PET. Additionally, reference tracing of systematic reviews and meta-analyses (on the topics of impulsivity more broadly), as well as response inhibition and delay discounting, specifically was done. The search terms were selected in keeping with the ‘pure measures’ of impulsivity within each dimension as suggested by Strickland et al. (2020). For DCS, these were: “delay discounting”, “temporal discounting”, “delayed reward” as well as each of the keywords separately. The database for response inhibition studies using the go/nogo and stop-signal paradigms in adults was obtained from a recent meta-analysis by Cieslik et al. (2023). We enriched this database by adding studies with adolescent participants for which we used the same search terms as presented in Cieslik et al. (2023), namely: “stop signal task”, “go no-go task”, “go nogo task”, “response inhibition”, “inhibition”, “action withholding”, “action cancellation”, “action inhibition”, “motor inhibition” and “inhibitory control”.

We included only results from peer-reviewed fMRI or perfusion PET experiments reporting results of whole-brain group analyses as coordinates in a standard neuroanatomical reference space (Talairach/Tournoux or Montreal Neurological Institute). Results from region-of-interest (ROI) analyses and studies with partial brain coverage were excluded. Only data from healthy participants (including healthy control groups from patient studies) with mean age >= 12 (with an absolute minimum age of individual participants no lower than 10) were retained. Studies with pharmacological interventions, connectivity-based analyses and single-subject reports were excluded. For studies reporting more than one eligible experiment obtained in the same sample, the reported coordinates were pooled to form a single experiment when included in the same meta-analysis (i.e. coordinates from go/nogo and stop-signal tasks in the same subject group were pooled). If each experiment included a different set of participants, coordinates were not pooled. In cases where different studies or experiments reported results from partly overlapping samples such as in Kable et al. (2007) and Kable et al. (2010), coordinates were pooled to form a single experiment and the smaller sample size of the two original experiments was used as the input to the analysis. In cases where any of the above criteria were unclear from screened publications, the corresponding authors were contacted. Lastly, authors of clinical studies that passed our inclusion criteria but reported pooled activation for clinical and healthy control groups were contacted for data from the healthy control group only. Of these, three authors responded and are indicated in the table of included studies in the supplementary material For a reporting checklist detailing analysis and study selection choices as suggested by Müller et al. (2018), see supplementary table S1.

Our contrasts of interest were, in general, analyses contrasting impulsive with non-impulsive, ‘controlled’ behaviour and vice versa, as impulsivity in pertinent paradigms is behaviourally expressed by a higher frequency of ‘impulsive responding’ such as commission errors (failure to inhibit action when necessary) or choices of smaller but sooner rewards (over larger but later ones). To differentiate the two types of contrasts, we refer to contrasts reflecting impulsive behaviour as ‘impulsive responding’ and to the reverse contrasts reflecting non-impulsive behaviour as ‘controlled responding’. Experiments reporting relative deactivations were interpreted as results of the opposite contrast to that specified (e.g., deactivation observed in a smaller sooner > larger later rewards contrast was interpreted as activation associated with larger later > smaller sooner rewards) unless otherwise specified in the respective publication. A detailed description of the selected contrasts for each impulsivity dimension is provided below. After the exclusion of unsuitable studies (see Fig. 1), the final sample consisted of 46 studies reporting 47 experiments on delayed-consequence sensitivity (21 reporting impulsive, 26 controlled responding and 24 subjective value) and 101 studies reporting 104 experiments on response inhibition (26 reporting impulsive and 96 controlled responding). Details on all studies included can be found in the supplementary material.

##### 1. Delayed-consequence sensitivity

Experiments were separated into 3 categories: impulsive responding, controlled responding and subjective value and separate meta-analyses were calculated for each category. For impulsive responding, results of smaller sooner (SS) > larger later (LL) rewards, immediate > delayed choice, and *β* > *δ* contrasts were selected, while for controlled responding the opposite contrasts were included, namely: LL > SS, delay > immediate and *δ* > *β*. *β* is theorised to reflect an ‘impatient system’ and is usually coded in fMRI paradigms as blocks of trials where immediate rewards are possible, while *δ* represents the ‘patient system’ and is coded as blocks of choices where only delayed choices occur (Laibson, 1997; McClure et al., 2004). We further included contrasts that tested for across-participant correlations between brain activity and the temporal discount parameter *k* (or similar constructs reflecting the degree to which individuals discount future rewards). As higher *k* indicates stronger impulsive tendencies, positive correlations were included in the meta-analysis of impulsive responding and negative correlations in the analysis of controlled responding. Lastly, choices between SS and LL rewards are highly influenced by the perceived subjective value of the rewards, which is believed to track the valuation processes during delay discounting tasks (Kable & Glimcher, 2007; Schüller et al., 2019). Therefore, parametric modulation and correlation of activity with subjective value were coded as a third category of experiments.

##### 2. Response Inhibition

Following the guidelines for performing well-powered fMRI meta-analyses (Eickhoff et al., 2016; Müller et al., 2018), we were not able to find a suitable amount of experiments reporting results of the direct comparison between impulsive and controlled responding (with only 15 for impulsive > controlled and 7 experiments for controlled > impulsive). We, therefore, selected experiments contrasting against control conditions not reflecting impulsivity like ‘Go’ conditions (no need for inhibition) or rest/fixation and then calculated the contrast of interest (impulsive vs. controlled) on the meta-analytical level. In particular, experiments that contrasted brain activation during commission errors or successful inhibition against baseline (Go, fixation or rest) were included. First, we calculated separate meta-analyses for impulsive responding > baseline and controlled responding > baseline, respectively. Next, we compared impulsive and controlled responding by calculating meta-analytic contrasts and conjunction analyses (for further details, see section Activation Likelihood Estimation below).

#### Activation likelihood estimation (ALE)

All meta-analyses were performed using the ALE algorithm for coordinate-based meta-analysis of neuroimaging studies (Turkeltaub et al., 2002; Eickhoff et al., 2009, 2012) that was implemented using in-house Matlab (version 2017a) tools. The analyses were executed as described previously (Kogler, Müller, Werminghausen, Eickhoff, & Derntl, 2020) and according to the best-practice guidelines for neuroimaging meta-analyses (Müller et al., 2018). The ALE algorithm aims to identify brain areas where activity across many experiments converges more strongly than would be expected from a random spatial association. Briefly, to reflect the spatial uncertainty of activations, each activation focus was modelled as a centre of a 3D Gaussian probability distribution based on empirical data of between-template and between-subject variance. The between-subject variance was weighted by the number of participants in the respective experiment. For a given experiment, the probability distributions of each focus were then combined and a union over all experiments‘ activation maps were computed. This yielded a voxelwise estimated activation likelihood map (i.e. a map of ALE scores), which describes the degree of spatial convergence across all experiments. Lastly, in order to identify ‘true‘ convergence, the ALE scores were compared to an analytically derived null distribution (Eickhoff et al., 2012) reflecting random spatial associations between activation maps for all experiments. Results were thresholded at p < .05 (family-wise error-corrected at cluster level with voxel-level cluster inclusion threshold at p < .001; Eickhoff et al., 2016).

#### Meta-analytic contrast and conjunction analyses

Contrast and conjunction analyses were calculated between meta-analytic results within each behavioural dimension (i.e. for impulsive vs. controlled) to directly compare impulsive and controlled responding for response inhibition and simplify peak extraction (see below). Commonalities between the two meta-analyses were assessed via conjunction analysis, which identifies voxels with significant convergence in both meta-analyses, calculated as the intersection of the cFWE-thresholded result maps. A cluster extent threshold of at least 5 voxels was applied to the resulting conjunction maps.

For contrast analyses, the voxel-wise differences between ALE scores of two meta-analyses were calculated and compared to a null distribution of difference scores. This null distribution was derived by pooling all experiments from the two meta-analyses and randomly dividing them into two groups of the same sample size as the original sets. This procedure was repeated 25,000 times to yield an empirical null distribution of ALE-score differences which the observed difference in ALE scores was tested against. The resulting voxel-wise nonparametric p values were thresholded at p < 0.05, cluster extent threshold of at least 5 voxels. While for delayed consequence sensitivity the number of included experiments was quite similar, for response inhibition the meta-analyses of controlled versus impulsive responding were unbalanced (96 vs 26 experiments). To accommodate for the higher power of the controlled responding > baseline meta-analysis, we employed a subsampling procedure described in detail in the supplementary methods.

#### Peak extraction

Next, we created a network comprising all regions involved in response inhibition (impulsive and controlled responding) and DCS (impulsive responding and subjective value). We thus combined the peaks of all meta-analytical networks into one single network. Peaks were extracted from the conjunction and contrast analyses between the meta-analyses of each behavioural task dimension (as described above). Thus, the peaks were on the one hand based on those regions that were found to be involved in more than one meta-analysis as well as those that showed stronger convergence in one compared to another meta-analysis (within the DCS and response inhibition domain, respectively). Peaks that lay in grey matter were thus extracted from the respective conjunction and contrast maps using fsl5 (Smith et al., 2004) [cluster] command with the minimum distance between peaks set to 15 mm. For peaks coming from different maps (for example conjunction and contrast maps) that were less than 15 mm apart from each other, we included only the peak with the higher z-score (Nostro et al., 2018). The extracted meta-analytic nodes and all result maps are available in the ANIMA database (Reid et al., 2016): https://anima.fz-juelich.de/studies/Gell_Impulsivity_2023

### Follow-up connectivity analyses for network characterisation

#### Participants

For all connectivity modelling, two different samples were used: one served as the discovery sample and one for replicating results. The discovery sample was chosen based on its cross-sectional design reflecting the sample used in the meta-analysis. This allowed us to use the same cut-off age range of 10 to 75 years present in the meta-analysis.

#### Discovery sample

Resting-state and anatomical (f)MRI data of 608 healthy subjects (395 female) aged 10-75 years were obtained from the extended Nathan Kline Institute dataset (Nooner et al., 2012). Only data from participants who had completed the full 10 min of scanning without excessive movement (defined here as mean framewise displacement of ≤ 0.5 mm) were included in further analyses, resulting in a final sample of n = 528 healthy subjects (338 female, age: 10-75 years). We used whole-brain T1 anatomical MPRAGE images (TR = 1900 ms; 1 mm isotropic voxels) and resting-state fMRI (rsfMRI) multiband echo-planar imaging (EPI) scans (TR = 1400 ms; 2 mm isotropic voxels; duration = 10 minutes; 440 volumes), acquired on a 3-T Siemens Magnetom scanner.

#### Replication sample

For replication, the minimally preprocessed data of a sample of unrelated healthy subjects (n = 339, 184 female, aged 22-35 years) were obtained from the full release of the Human Connectome Project dataset (Van Essen et al., 2013). We excluded participants with incomplete resting-state scans or excessive movement (mean framewise displacement of > 0.2 mm as used previously by e.g., Yang et al., 2016) resulting in a final sample of n = 336 subjects (183 female, age: 22-35 years). The rsfMRI HCP scanning protocol involved acquiring whole-brain multiband gradient-echo EPI volumes on a 3-T Siemens “Connectome Skyra” scanner (TR = 720 ms, 2 mm isotropic voxels). Four rsfMRI sessions with 1,200 volumes in total (14 min and 24 s) were acquired over two consecutive days, with one left-to-right (LR) and one right-to-left (RL) encoding direction acquired on each day. For the purposes of replicating our findings based on the eNKI sample, only data from the first session on the first day was used (so-called “rest1LR”).

#### Preprocessing

The eNKI data were preprocessed using fMRIPrep version 20.1.1 (Esteban et al., 2018; fMRIPrep 2020), which is based on Nipype version 1.5.0 (Gorgolewski et al., 2011; Nipype 2017). For a detailed description of each step, see supplementary methods. Briefly, this included skull-stripping, head-motion correction and slice-time correction. The BOLD images were then coregistered to the native space of the subjects’ T1w image, normalised to MNI space and motion-corrected.

The HCP data used here were minimally preprocessed. The preprocessing pipeline has been described in detail elsewhere (Glasser et al., 2016). Briefly, this included gradient distortion correction, image distortion correction, registration to subjects’ T1w image and to MNI standard space followed by intensity normalisation of the acquired rsfMRI images, and ICA FIX denoising (Salimi-Khorshidi et al., 2014).

For both datasets, additional denoising steps were undertaken using fMRIPrep output files or data provided by the HCP and in-house scripts in MATLAB (version 2019b). First, we regressed mean time courses of 2 tissue classes (white matter and cerebrospinal fluid) and the global signal which has been shown to reduce motion-related artefacts (Ciric et al., 2017). Next, data were linearly detrended, bandpass-filtered at 0.01 – 0.1 Hz and spatially smoothed using a Gaussian kernel of FWHM = 5 mm.

#### Community detection and network measures

After averaging the time series from all grey-matter voxels within 5-mm spheres around the meta-analytically derived coordinates, node-to-node functional connectivity was calculated as the Pearson correlation between the time courses of each node. The resulting connectivity matrix for each participant was z-scored using Fisher’s-z transformation and averaged across all participants. We employed the Louvain algorithm (Blondel et al., 2008), a stochastic method, for identifying distinct communities within a network by optimising Q, a modularity score (Betzel, 2020). For this, we used the community_louvain.m function from the Matlab-based Brain Connectivity Toolbox (Rubinov & Sporns, 2010). The averaged connectivity matrix between all meta-analytic nodes was used as the input. We finetuned the community assignment by using the communities resulting from applying the algorithm to the connectivity matrix as an additional input and repeated the procedure until Q remained constant. Given the greedy stochastic nature of the algorithm (Good, de Montjoye, & Clauset, 2010), community assignment was evaluated by repeating the procedure 1000 times to obtain an agreement matrix. To evaluate the node roles in the final community partition, we calculated the participation coefficient (participation_coef_sign.m) and within-module degree z-score (module_degree_zscore.m). The participation coefficient identifies if a node‘s connections are distributed across communities or clustered within a community and reflects between-module integration at high values and segregation at low values. The within-module degree z-score describes the connectedness of a node to its own community relative to other nodes in the same community and thus reflects within-module integration.

#### Seed-voxel connectivity gradients

While the above-described community detection identifies communities based on node-to-node connectivity profiles, we additionally investigated network organisation based on the ‘node-to-rest of the brain’ connectivity profiles (i.e. seed-to-voxel correlations). In order to determine if nodes located in the same community displayed similar connectivity to the rest of the brain, we identified principal axes of variation in the connectivity profiles across all nodes. This technique was recently used to determine spatial variation in both node-to-node (Margulies et al., 2016) and seed-to-voxel (J. Zhang et al., 2019) connectivity, as well as structural characteristics such as microstructure (Paquola et al., 2019) across the cortex. Seed-to-voxel connectivity was calculated as Pearson correlation between the mean time courses of each node and all remaining grey-matter voxels in the brain, resulting in one connectivity map for each node per subject. Maps for each node were Fisher Z-transformed before averaging across participants. Next, we constructed a node-by-node similarity matrix, by transforming the averaged (3D) seed-to-voxel connectivity map of each node into a vector and correlating the resulting vectors from each node (resulting in a 21x21 matrix). To this matrix, we then applied principal component analysis using the BrainSpace toolbox (Vos de Wael). Only the top 20% of node similarities were retained (i.e. sparsity parameter). The remaining parameters were kept the same as in previous work by Marguilles et al. (2016), with α set to 0.05. We repeated the gradient decomposition using diffusion map embedding (Coifman et al., 2005) and varying levels of sparsity (30% and 40%) in order to confirm our results were not subject to the choice of dimensionality reduction algorithm or parameters.

### PET-based receptor density analysis

To investigate the relationship between neurotransmitter receptor/transporter density and community organisation, we used PET-derived whole-brain maps available in the JuSpace toolbox (Dukart et al., 2021) available online (https://www.fz-juelich.de/inm/inm-7/EN/Resources/_doc/JuSpace.html?nn=2463520). For the analysis, we only used receptor and transporter maps for neurotransmitters theoretically related to impulsivity: serotonin, norepinephrine and dopamine (Chamberlain & Sahakian, 2007; Dalley et al., 2011; Dalley & Robbins, 2017). In particular, for serotonin, we utilised the 5HT1a, 5HT1b, 5HT2a and serotonin transporter maps (SERT) (Savli et al., 2012), norepinephrine transporter map NAT (Hesse et al., 2017) for norepinephrine, and D1 (Kaller et al., 2017), D2 (Alakurtti et al., 2015) and dopamine transporter maps (Dukart et al., 2018) for dopamine. All PET maps were acquired from healthy volunteers and rescaled to a minimum of 0 and a maximum of 100, for further details, see Dukart et al. (2021).

First, all the above PET maps were resampled from 3 mm isotropic voxels to 2 mm isotropic voxels using the fsl5 [flirt] command. For each node, we then averaged the receptor density values in all grey matter voxels within 5 mm diameter spheres around each coordinate. Next, the node-wise receptor density was correlated (using Spearman rank correlation) with within-module degree z-score and participation coefficient derived from the community organisation. Correlations that displayed at least moderate effect size (>+-0.3) in both our discovery and replication datasets were then tested against a spatially informed null model for significance using permutation testing. To this end, we created 1000 random networks by randomly sampling coordinates from a conservative grey-matter mask. To mirror the spatial properties of our impulsivity network in the randomly sampled networks, we restricted the minimum, mean and maximum Euclidean distance between the sampled nodes to be within 1 standard deviation from the impulsivity network‘s minimum, mean and maximum values, respectively. We then calculated Spearman rank correlation between receptor density in nodes of each random network and our empirically derived measures of integration and segregation to estimate a null distribution. The empirical rank correlation was then compared to the estimated null. Correlation coefficients higher than 95% of the random correlations were interpreted as significant.

## Supporting information

Included studies

Supplementary results

