## Supplementary material for "Charting the brain networks of impulsivity: Meta-analytic synthesis, functional connectivity modelling and neurotransmitter associations": Included studies

### Included Studies for Delay Consequence Sensitivity

| Impulsive Decision-making | | | | | |
|---|---|---|---|---|---|
| Author | N | Contrast | Space | Were in-scanner presented rewards adapted based on subjective value? | Source (of coordinates) |
| Albrecht et al., 2011 | 28 | immediate > delay | TAL | not subject adapted | Table S4 |
| Albrecht et al., 2013 | 30 | now > later | TAL | not subject adapted | Table S1 |
| Banich et al., 2013 | 29 | now > later | MNI | not subject adapted | paper |
| Christakou et al., 2011 | 40 | immediate > delay | TAL | subject adapted | paper |
| de Water et al., 2017 | 58 | correlation of sensitivity to immediate reward in immediate > delay | MNI | not subject adapted | paper |
| Deshpande et al., 2019 | 54 | immediate and delay > delay and delay | MNI | subject adapted | paper |
| Elton et al., 2017 | 95 | positive correlation with impulsive choice ration (similar to k) in immediate & delay > control | MNI | not subject adapted | paper |
| Eppinger et al., 2012 | 30 | beta (immediate & delay > all choice options) | TAL | not subject adapted | paper |
| Faralla et al., 2015 | 28 | (immediate > delay) > control | MNI | not subject adapted | Table 5 |
| Hamilton et al. 2020 | 27 | SS > LL | MNI | subject adapted | neurovault |
| Kable and Glimcher 2010 | 22 | immediate and delay > delay and delay | MNI | subject adapted | paper |

| | | | | | |
|---|---|---|---|---|---|
| Luo et al., 2009 | 37 | immediate > delay | MNI | subject adapted | paper |
| Mavrogiorgou et al., 2016 | 20 | immediate > delay | MNI | subject adapted | paper |
| McClure et al., 2004 | 14 | beta (immediate & 0.5*delay) | MNI | not subject adapted | paper |
| Norman et al., 2017 | 20 | immediate > delay | TAL | subject adapted | paper |
| Pine et al., 2009 | 24 | correlation with discount factor D | MNI | not subject adapted | paper |
| Samanez-Larkin et al., 2011 | 25 | beta | TAL | not subject adapted | paper |
| Sripada et al., 2011 | 20 | immediate and delay > delay and delay | MNI | not subject adapted | paper |
| Wittmann et al., 2010 | 13 | immediate > delay | TAL | not subject adapted | table S7 |
| Xu et al., 2009 | 18 | immediate and delay > delay and delay | MNI | not subject adapted | table S2 |
| Zhuang et al., 2020 | 16 | sooner > delayed (main effect of delay decision) | MNI | not subject adapted | paper |

| Controlled Decision-making | | | | | |
|---|---|---|---|---|---|
| Author | N | Contrast | Space | Were in-scanner presented rewards adapted based on subjective value? | Source (of coordinates) |
| Banich et al., 2013 | 29 | later > now | MNI | not subject adapted | paper |
| Christakou et al., 2011 | 40 | delay > immediate | TAL | subject adapted | paper |
| de Water et al., 2017 | 58 | delay > immediate | MNI | not subject adapted | paper |
| Deshpande et al., 2019 | 54 | delay and delay > immediate and delay | MNI | subject adapted | paper |
| Elton et al., 2017 | 95 | neg corr. with impulsive choice ratio | MNI | not subject adapted | paper |
| Eppinger et al., 2012 | 30 | delta (delay & immediate) | TAL | not subject adapted | paper |
| Faralla et al., 2015 | 28 | (delay > immediate) > control | MNI | not subject adapted | Table 6 |
| Hamilton et al. 2020 | 27 | LL > SS | MNI | subject adapted | neurovault |
| Hill et al., 2017 | 25 | delay > immediate | MNI | subject adapted | Table S2 |
| Kable and Glimcher 2007 | 10 | delay > immediate | TAL | subject adapted | Table S5 |
| Kable and Glimcher 2010 | 22 | delay and delay > immediate and delay | TAL | subject adapted | paper |
| King et al., 2016 | 36 | LL > SS | MNI | subject adapted | authors |
| Laube et al., 2020 | 48 | LL > SS | MNI | subject adapted | neurovault |
| Luo et al., 2009 | 37 | delay > immediate | MNI | subject adapted | paper |

| | | | | | |
|---|---|---|---|---|---|
| Luo et al., 2012 | 21 | LL > SS | MNI | subject adapted | paper |
| McClure et al., 2004 | 14 | delta (delay & immediate > baseline) | MNI | not subject adapted | paper |
| Miedl et al., 2015 | 15 | delay > immediate | TAL | not subject adapted | paper |
| Norman et al., 2017 | 20 | delay > immediate | TAL | subject adapted | paper |
| O'Connell et al., 2018 | 26 | delay > immediate | MNI | subject adapted | paper |
| Samanez-Larkin et al., 2011 | 25 | delta | TAL | not subject adapted | paper |
| Sripada et al., 2011 | 20 | delay > immediate | MNI | not subject adapted | paper |
| Van den Bos et al., 2014 | 22 | LL > SS | MNI | subject adapted | Table S1 |
| Waegeman et al., 2014 | 41 | delay > immediate | MNI | not subject adapted | paper |
| Wang et al., 2017 | 21 | delay > immediate | MNI | not subject adapted | paper |
| Wittmann et al., 2007 | 13 | delay > immediate | TAL | not subject adapted | paper |
| Wittmann et al., 2010 | 13 | delay > immediate | TAL | not subject adapted | Table S7 |
| Zhuang et al., 2020 | 16 | delayed > sooner (main effect of delay decision) | MNI | not subject adapted | paper |

| Subjective value | | | | | |
|---|---|---|---|---|---|
| Author | N | Contrast | Space | Were in-scanner presented rewards adapted based on subjective value? | Source (of coordinates) |
| Castrellon et al., 2019 | 21 | subjective value (parametric modulation) | MNI | not subject adapted | paper |
| Cox and Kable 2014 | 20 | subjective value (parametric modulation) | MNI | subject adapted | paper |
| Eppinger et al., 2017 | 50 | correlation with subjective value | TAL | subject adapted | paper |
| Hare et al., 2014 | 25 | correlation with discounted stimulus value (SV) | MNI | not subject adapted | paper |
| Jimura et al., 2013 | 43 | correlation with subjective value | TAL | subject adapted | paper |
| Kable and Glimcher 2007 | 10 | correlation with subjective value, discount rate | TAL | subject adapted | Table S5 |
| Kable and Glimcher 2010 | 22 | correlation with subjective value | TAL | subject adapted | paper |
| King et al., 2016 | 36 | subjective value (parametric modulation) | MNI | subject adapted | authors |
| Lempert et al., 2017 | 35 | subjective value (parametric modulation) | MNI | not subject adapted | paper |
| Liu et al., 2012 | 19 | correlation with subjective value | TAL | subject adapted | paper |
| Luo et al., 2012 | 21 | subjective value, stochasticity, discounting factor (k) | MNI | subject adapted | paper |
| Massar et al., 2015 | 23 | correlation with subjective value | TAL | subject adapted | paper |
| Murawski, et al., 2012 | 13 | subjective value > baseline | MNI | subject adapted | Table S5 |

| | | | | | |
|---|---|---|---|---|---|
| O'Connell et al., 2018 | 26 | subjective value (parametric modulation) | MNI | subject adapted | paper |
| Peters and Büchel 2009 | 22 | correlation with subjective value | MNI | subject adapted | paper |
| Peters and Büchel 2010 | 30 | correlation with subjective value | MNI | subject adapted | In text |
| Pine et al., 2009 | 24 | correlation with subjective value (discounted utility) | MNI | not subject adapted | paper |
| Prevost et al., 2010 | 18 | correlation with subjective value | MNI | subject adapted (force) | paper |
| Ripke et al., 2012 | 27 | subjective value (parametric modulation) | MNI | subject adapted | paper |
| Samanez-Larkin et al., 2011 | 25 | correlation with subjective value | TAL | not subject adapted | paper |
| Sasse et al., 2017 | 22 | subjective value (parametric modulation) | MNI | subject adapted | paper |
| Seaman et al., 2018 | 75 | correlation with subjective value | MNI | not subject adapted | paper |
| Sripada et al., 2011 | 20 | correlation with subjective value | MNI | not subject adapted | paper |
| Wang et al., 2014 | 28 | subjective relative value, corr. with value of immediate option | MNI | not subject adapted (adapted from pilot study matching in age and gender) | paper |
| Wiehler et al., 2017 | 23 | subjective value (parametric modulation) | MNI | subject adapted | authors |

### Included Studies for Response Inhibition

| Successful Inhibition Database | | | | | |
|---|---|---|---|---|---|
| Author | n | contrast | space | paradigm | type of error |
| Aron and Poldrack, 2006 | 13 | Correct Stop > Correct Go | MNI | stop signal task | SM Table 2 |
| Berkmann et al., 2014 | 60 | Correct Stop > Correct Go | MNI | stop signal task | Table 2 |
| Bobb et al., 2011 | 13 | Correct Stop > Correct Go | MNI | stop signal task | Table 2 |
| Boecker et al., 2011 | 15 | Correct Stop > Correct Go | MNI* | stop signal task | SM Table 1 |
| Boehler et al., 2010 | 15 | Correct Stop > Go | MNI | stop signal task | Table 3 |
| Cai and Leung, 2009 | 12 | Correct Stop > Correct Go | MNI | stop signal task | Table 1 |
| Cai and Leung, 2011 | 23 | Correct Stop > Go | MNI | stop signal task | SM Table 1 |
| Cai et al. 2014 | 19 | Correct Stop > Correct Go | MNI | stop signal task | Table 2 |
| Chen et al., 2015 | 25 | Correct NoGo > Go | MNI | go/no-go | Table 2 |
| Chevrier et al., 2007 | 14 | correct stop > go | TAL | stop signal task | Table 1 |
| Chikara et al., 2018 | 20 | Correct Stop > Correct Go | MNI | stop signal task | Table 3A |
| Chikazoe et al., 2009a | 22 | Correct Stop > Correct uncertain go trials | MNI | stop signal task | Table 2 |
| Chikazoe et al., 2009b | 25 | correct No-Go > infrequent correct Go | MNI | go/no-go | Table 1 |
| | | | | go/no-go | Table 2 |

| | | | | | |
|---|---|---|---|---|---|
| Congdon et al., 2014 | 62 | Correct Stop > Correct Go | MNI | stop signal task | Table 2 |
| Coxon et al., 2016_1 | 20 | Correct Stop > Correct Go | MNI | stop signal task | Table 2 (young) |
| Coxon et al., 2016_2 | 20 | Correct Stop > Correct Go | MNI | stop signal task | Table 2 (old) |
| Czapla et al., 2017 | 21 | Correct NoGo > Correct Go | MNI | go/no-go | SM Table 4 |
| | | | | go/no-go | SM Table 5 |
| Dambacher et al., 2015 | 15 | Correct NoGo > Go | TAL | go/no-go | Table 1 |
| Fassbender et al., 2004 | 18 | Correct Inhibitions > tonic activation | TAL | go/no-go | Table 2 |
| Fauth-Bühler et al., 2012 | 18 | successful stop > baseline | MNI | stop signal task | from authors |
| Fedota et al., 2015 | 16 | Correct NoGo > Correct Go | MNI | go/no-go | Table 1 |
| Fuentes-Claramonte et al., 2016 | 57 | Correct NoGo > Correct frequent Go | MNI | go/no-go | Table 2 |
| | | Correct NoGo > Correct Infrequent Go | | go/no-go | Table 2 |
| Ganos et al., 2014 | 15 | Correct Stop > Go | MNI | stop signal task | SM Table 2 |
| Garavan et al., 1999 | 14 | correct Inhibition > active baseline | TAL | go/no-go | Table 1 |
| Garavan et al., 2002 | 14 | correct Inhibition > active baseline | TAL | go/no-go | Table 1 |
| Garavan et al., 2003 | 16 | correct Inhibition > active baseline | TAL | go/no-go | Table 1 |
| Geng et al., 2009 | 16 | Correct NoGo > Correct Go | MNI | go/no-go | Table 3 |
| Ghahremani et al., 2012 | 18 | Correct Stop > Correct Go | MNI | stop signal task | Table 3 |

| | | | | | |
|---|---|---|---|---|---|
| Harle et al., 2016 | 34 | Correct Stop > Correct Go | TAL | stop signal task | SM Table 2 |
| Hester et al., 2004a | 15 | successful inhibition vs. Baseline | TAL | go/no-go | Table 1 |
| Hester et al., 2004b | 15 | Successful inhibition vs go | TAL | go/no-go | Table 3 |
| Hough et al., 2016 | 22 | Correct NoGo > Correct Go | MNI | go/no-go | SM Table |
| Hsu et al., 2017 | 20 | Correct NoGo > Go | MNI | go/no-go | Table 2 |
| Hughes et al., 2012 | 10 | Correct stop > baseline | MNI | stop signal task | Table 4 |
| Hughes et al., 2013 | 15 | Correct stop > baseline | MNI | stop signal task | Table 3 |
| Jahfari et al., 2011 | 20 | Correct Stop > Go | MNI | stop signal task | Table 4 |
| Jahfari et al., 2012 | 16 | Correct Stop > Go | MNI | stop signal task | Table 5 |
| Jahfari et al., 2015 | 23 | Correct Stop > Go | MNI | stop signal task | Table 3 |
| Kaladjian et al., 2007 | 21 | correct No-Go > Correct Go | TAL | go/no-go | Table 3 |
| Kaladjian et al., 2009a | 20 | correct No-Go > Correct Go | TAL | go/no-go | Table 3 |
| Kaladjian et al., 2009b | 10 | correct No-Go > Correct Go | TAL | go/no-go | Table 3 |
| | | correct No-Go > Correct Go | | go/no-go | Table 3 |
| Kelly et al., 2004 | 15 | Successful inhibition vs baseline | TAL | go/no-go | Table1 |
| Kenner et al., 2010 | 24 | Correct Stop > Correct Go | MNI | stop signal task | SM Table 3 |
| Kiehl et al., 2000 | 14 | correct Inhibition > active baseline | MNI | go/no-go | Table 1 |
| Ko et al., 2014 | 23 | Correct NoGo > Go | MNI | go/no-go | Table 2 |

| | | | | | |
|---|---|---|---|---|---|
| Ko et al., 2016 | 32 | Correct Stop > Correct Go | MNI | stop signal task | Table 1A |
| | | Correct Stop > Correct Go | | stop signal task | Table 1B |
| Köhler et al., 2018 | 33 | Correct NoGo > Correct Go | MNI | go/no-go | SM Table 1 |
| Lavallee et al., 2014 | 21 | Correct Stop > Go | MNI | stop signal task | Table 2 |
| Liddle et al. , 2001 | 16 | correctNoGo > correctGo | MNI | go/no-go | Table 3 |
| Lorenz et al., 2015 | 38 | Corect Stop > Go | MNI | stop signal task | SM Table 1 |
| Marco-Pallarés et al., 2008 | 10 | correct Stop > correct Go | MNI | stop signal task | Table 1 |
| Mazzola-Pomietto et al., 2009 | 16 | correct No-Go > Correct Go | TAL | go/no-go | Table 3 |
| Mohammadi et al., 2015 | 17 | Correct Stop > Go | MNI | stop signal task | SM Table 2 |
| Montojo et al., 2013 | 30 | Correct Stop > Correct Go | MNI | stop signal task | Table 2 |
| Nakata et al., 2008 | 15 | Correct NoGo > Correct Go | TAL | go/no-go | Table 5 |
| O'Connor et al., 2012 | 18 | Correct NoGo > active baseline | MNI | go/no-go | Table 1 |
| Rae et al., 2014 | 17 | Stop specified correct > go specified correct | MNI | stop signal task | SM Table 2 |
| | | Stop select correct > go select correct | | stop signal task | SM Table 2 |
| Rodriguez-Pujadas et al., 2014 | 33 | Correct Stop > Correct Go | MNI* | stop signal task | Table 2 |
| Rothmayr et al., 2011 | 12 | correct No-Go > Correct Go | MNI | go/no-go | Table 2 |
| Rubia et al., 2006_1 | 21 | successful NoGo > successful Go | TAL | go/no-go | Table 2 |

| | | | | | |
|---|---|---|---|---|---|
| Rubia et al., 2006_2 | 25 | successful no-go > successful go | TAL | go/no-go | Table 2 |
| Schel et al., 2014 | 24 | Correct Stop > Correct Go | MNI | stop signal task | Table 4 |
| Sebastian et al., 2012 | 24 | Correct NoGo > Correct Go | MNI | go/no-go | Table 5 |
| | | Correct Stop > Correct Go | | stop signal task | Table 5 |
| Sebastian et al., 2013a | 48 | Correct NoGo > active baseline including go | MNI | go/no-go | SM Table 1 |
| | | Correct Stop > Correct Go | | stop signal task | SM Table 1 |
| Sebastian et al., 2013b_1 | 24 | Correct NoGo > active baseline including go | MNI | go/no-go | Table 3 |
| | | Correct Stop versus correct Go | | stop signal task | Table 3 |
| Sebastian et al., 2013b_2 | 21 | correct NoGo > correct congruent go | MNI | go/no-go_hybrid response interference | Table 2 |
| | | Correct Stop versus correct congruent Go | | stop signal task_hybrid response interference | Table 2 |
| Sebastian et al., 2017 | 80 | Correct Stop > Correct Go | MNI | stop signal task | from authors |
| Sharp et al., 2010 | 26 | Correct Stop > Correct Go | MNI | stop signal task | SM Table 1 |
| Steele et al. 2014 | 102 | Correct NoGo > Correct Go | MNI | go/no-go | Table 2 |
| Swann et al. 2012 | 16 | Correct Stop > Correct Go | MNI | stop signal task | SM Table 2 |
| Tabu et al., 2011 | 13 | Correct Stop > Correct Go | MNI | stop signal task | text, page 280 |

| | | | | | |
|---|---|---|---|---|---|
| Tabu et al., 2012 | 13 | Correct Stop > Correct Go | MNI | stop signal task | SM Table 1 (hand task) |
| | | Correct Stop > Correct Go | | stop signal task | SM Table 1 (foot task) |
| Van der Meer et al., 2013 | 19 | Correct Stop > Correct Go | MNI | stop signal task | SM Table 3A |
| Van Eijk et al., 2015_1 | 18 | Correct NoGo > Correct Go | MNI | go/no-go | Table 4 |
| | | Correct Stop > Correct Go | | stop signal task | Table 4 |
| van Eijk et al., 2015_2 | 25 | Correct NoGo > Correct congruent Go | MNI | go/no-go_hybrid response interference | Table 5 |
| | | Correct stop > correct congruent go | | stop signal task_hybrid response interference | Table 5 |
| Walther et al., 2010 | 17 | correct No-Go > Correct Go | MNI | go/no-go | Table 1 |
| Wilbertz et al., 2014 | 49 | Correct Stop > Go | MNI | stop signal task | SM Table 2 |
| Xu et al., 2015 | 18 | Correct Stop > Go | MNI | stop signal task | Table 2 |
| Xu et al., 2017 | 21 | Correct Stop > Correct Go | MNI* | stop signal task | SM Table 1 |
| Xue et al., 2008 | 15 | Correct Stop > Correct Go | MNI | stop signal task | SM Table 1 (manual) |
| | | Correct Stop > Correct Go | | stop signal task | SM Table 1 (letter naming) |

| | | | | | |
|---|---|---|---|---|---|
| Zandbelt and Vink, 2010 | 24 | Correct Stop > Go | MNI | stop signal task | SM Table 3B |
| Zandbelt et al. 2011 | 22 | Correct Stop > Go | MNI | stop signal task | SM Table 11 |
| Zheng et al., 2008 | 18 | Correct Stop > Go | TAL | go/no-go | Table 1 |
| Lock et al., 2011 | 13 | correct No-Go > Correct Go | TAL | go/no-go | Table 2 |
| Nosarti et al., 2006 | 14 | (Correct no-go – go) – (correct 'odd' (aka go acting as a control for low freq of nogo) – go) | TAL | go/no-go | In text pg. 268 |
| Durston et al., 2006 | 11 | Successful No-go > Go | MNI | go/no-go | Table 2 |
| He et al., 2014 | 30 | Correct No-go > Go | MNI | go/no-go | Table 3 |
| Schulz et al., 2004 | 9 | Correct nogo > Correct Go | TAL | go/no-go | Table 2 |
| Lee et al., 2018 | 34 | Correct No-go > Go | MNI | go/no-go | Table 3 |
| Heitzeg et al., 2010 | 20 | Correct No-go > Go | MNI | go/no-go | Table 2 |
| Mulder et al., 2008 | 12 | Correct No-go > Go | MNI | go/no-go | Table 2 |
| White et al., 2014 | 1133 | stop success > baseline | MNI | stop signal task | SM Table 3 |
| Lim et al., 2015 | 27 | Stop Success > Go | MNI | stop signal task | SM Table |
| Bennett et al., 2009 | 11 | Correct No-go – Correct go | TAL | go/no-go | Table 2 |
| Cascio et al., 2015 | 37 | Correct No-Go > Correct Go | MNI | go/no-go | Table 3 |
| Rubia et al., 2013 | 66 | Stop Success > Go | TAL | stop signal task | Table 2 |

| Failure of Inhibition Database | | | | | |
|---|---|---|---|---|---|
| Author | N | contrast | space | paradigm | source of data |
| Boecker et al., 2011 | 15 | failed stop - Go | MNI | stop signal task | SM Table 2 |
| Boehler et al., 2010 | 15 | unsuccessful stop > go | MNI | stop signal task | SM Table 4 and 3 |
| Chen et al., 2015 | 25 | failed nogo - go | MNI | go/no-go | Table 2 |
| Chevrier et al., 2007 | 14 | unsuccessful stop vs baseline | TAL | stop signal task | Table 1 |
| Dambacher et al., 2015 | 15 | False alarm > go | TAL | go/no-go | Table 1 |
| Garavan et al., 2002 | 14 | error vs. Baseline | TAL | go/no-go | Table 1 |
| Garavan et al., 2003 | 16 | error vs. Baseline | TAL | go/no-go | Table 1 |
| Harle, 2016 | 34 | stop error > go | TAL | stop signal task | SM Table 3 |
| Hester et al., 2004 | 15 | error vs. Baseline | TAL | go/no-go | Table 2 |
| Hester et al., 2005 | 13 | error vs correct go | TAL | go/no-go | Table 1 |
| Hough et al., 2015 | 22 | unsuccessful inhibition vs baseline | MNI | go/no-go | SM Table 5 |
| Hsu et al., 2017 | 20 | unsuccessful inhibition vs go | MNI | go/no-go | Table 3 |
| Hughes et al., 2012 | 10 | stop failure vs baseline | MNI | stop signal task | Table 4 |
| Hughes et al., 2013 | 15 | stop failure vs baseline | MNI | stop signal task | Table 3 |

| | | | | | |
|---|---|---|---|---|---|
| Jahfari, 2011 | 20 | failed stop > go | MNI | stop signal task | Table 4 |
| Jahfari, 2012 | 16 | failed stop > go | MNI | stop signal task | Table 5 |
| Kiehl et al., 2000 | 14 | error vs baseline (error of comission) | MNI | go/no-go | Table 1 |
| Ko et al. 2014 | 23 | unsuccessful inhibition - go | MNI | go/no-go | Table 3 |
| Mohammadi et al., 2015 | 17 | unsuccessful stop > go | MNI | stop signal task | SM Table 3 |
| Rubia et al., 2003 | 20 | unsuccessful stop vs go | TAL | stop signal task | Table 1 |
| Sharp et al., 2010 | 26 | incorrect stop vs correct go | MNI | stop signal task | SM Table 4 |
| Steele et al., 2014 | 102 | error vs correct go | MNI | go/no-go | Table 1 and 3 |
| Xu et al., 2017 | 21 | failed stop > go | TAL | stop signal task | SM Table 2 |
| White et al., 2014 | 1133 | stop failure > baseline | MNI | stop signal task | SM Table 3 |
| Lim et al., 2015 | 27 | stop failure > go | MNI | stop signal task | SM Table |
| Halari et al., 2009 | 21 | stop failure – go | TAL | stop signal task | Table 3 |

### References: Delay Consequence Sensitivity


Albrecht, K., Volz, K. G., Sutter, M., & Cramon, D. Y. von. (2013). What Do I Want and When Do I Want It: Brain Correlates of Decisions Made for Self and Other. PLOS ONE, 8(8), e73531. doi: 10.1371/journal.pone.0073531

Albrecht, K., Volz, K. G., Sutter, M., Laibson, D. I., & von Cramon, D. Y. (2011). What is for me is not for you: Brain correlates of intertemporal choice for self and other. Social Cognitive and Affective Neuroscience, 6(2), 218–225. doi: 10.1093/scan/nsq046

Banich, M. T., De La Vega, A., Andrews-Hanna, J. R., Mackiewicz Seghete, K., Du, Y., & Claus, E. D. (2013). Developmental trends and individual differences in brain systems involved in intertemporal choice during adolescence. Psychology of Addictive Behaviors, 27(2), 416–430. doi: 10.1037/a0031991

Bos, W. van den, Rodriguez, C. A., Schweitzer, J. B., & McClure, S. M. (2014). Connectivity Strength of Dissociable Striatal Tracts Predict Individual Differences in Temporal Discounting. Journal of Neuroscience, 34(31), 10298–10310. doi: 10.1523/JNEUROSCI.4105-13.2014

Castrellon, J. J., Young, J. S., Dang, L. C., Cowan, R. L., Zald, D. H., & Samanez-Larkin, G. R. (2019). Mesolimbic dopamine D2 receptors and neural representations of subjective value. Scientific Reports, 9(1), 20229. doi: 10.1038/s41598-019-56858-1

Christakou, A., Brammer, M., & Rubia, K. (2011). Maturation of limbic corticostriatal activation and connectivity associated with developmental changes in temporal discounting. NeuroImage, 54(2), 1344–1354. doi: 10.1016/j.neuroimage.2010.08.067

Cox, K. M., & Kable, J. W. (2014). BOLD Subjective Value Signals Exhibit Robust Range Adaptation. Journal of Neuroscience, 34(49), 16533–16543. doi: 10.1523/JNEUROSCI.3927-14.2014

de Water, E., Mies, G. W., Figner, B., Yoncheva, Y., van den Bos, W., Castellanos, F. X., … Scheres, A. (2017). Neural mechanisms of individual differences in temporal discounting of monetary and primary rewards in adolescents. NeuroImage, 153, 198–210. doi: 10.1016/j.neuroimage.2017.04.013

Deshpande, H. U., Mellis, A. M., Lisinski, J. M., Stein, J. S., Koffarnus, M. N., Paluch, R., … Bickel, W. K. (2019). Reinforcer pathology: Common neural substrates for delay discounting and snack purchasing in prediabetics. Brain and Cognition, 132, 80–88. doi: 10.1016/j.bandc.2019.03.003

Elton, A., Smith, C. T., Parrish, M. H., & Boettiger, C. A. (2017). Neural Systems Underlying Individual Differences in Intertemporal Decision-making. Journal of Cognitive Neuroscience, 29(3), 467–479. doi: 10.1162/jocn_a_01069

Eppinger, B., Heekeren, H. R., & Li, S.-C. (2018). Age differences in the neural mechanisms of intertemporal choice under subjective decision conflict. Cerebral Cortex, 28(11), 3764–3774. doi: 10.1093/cercor/bhx239

Eppinger, B., Nystrom, L. E., & Cohen, J. D. (2012). Reduced Sensitivity to Immediate Reward during Decision-Making in Older than Younger Adults. PLOS ONE, 7(5), e36953. doi: 10.1371/journal.pone.0036953

Faralla, V., Benuzzi, F., Lui, F., Baraldi, P., Dimitri, N., & Nichelli, P. (2015). Neural correlates in intertemporal choice of gains and losses. Journal of Neuroscience, Psychology, and Economics, 8(1), 27–47. doi: 10.1037/npe0000032

Hamilton, K. R., Smith, J. F., Gonçalves, S. F., Nketia, J. A., Tasheuras, O. N., Yoon, M., … Shackman, A. J. (2020). Striatal bases of temporal discounting in early adolescents. Neuropsychologia, 144, 107492. doi: 10.1016/j.neuropsychologia.2020.107492

Hare, T., Hakimi, S., & Rangel, A. (2014). Activity in dlPFC and its effective connectivity to vmPFC are associated with temporal discounting. Frontiers in Neuroscience, 8. Retrieved from https://www.frontiersin.org/articles/10.3389/fnins.2014.00050

Hill, P. F., Yi, R., Spreng, R. N., & Diana, R. A. (2017). Neural congruence between intertemporal and interpersonal self-control: Evidence from delay and social discounting. NeuroImage, 162, 186–198. doi: 10.1016/j.neuroimage.2017.08.071

Jimura, K., Chushak, M. S., & Braver, T. S. (2013). Impulsivity and Self-Control during Intertemporal Decision Making Linked to the Neural Dynamics of Reward Value Representation. Journal of Neuroscience, 33(1), 344–357. doi: 10.1523/JNEUROSCI.0919-12.2013

Kable, J. W., & Glimcher, P. W. (2007). The neural correlates of subjective value during intertemporal choice. Nature Neuroscience, 10(12), 1625–1633. doi: 10.1038/nn2007

Kable, J. W., & Glimcher, P. W. (2010). An “As Soon As Possible” Effect in Human Intertemporal Decision Making: Behavioral Evidence and Neural Mechanisms. Journal of Neurophysiology, 103(5), 2513–2531. doi: 10.1152/jn.00177.2009

King, J. A., Geisler, D., Bernardoni, F., Ritschel, F., Böhm, I., Seidel, M., … Ehrlich, S. (2016). Altered Neural Efficiency of Decision Making During Temporal Reward Discounting in Anorexia Nervosa. Journal of the American Academy of Child & Adolescent Psychiatry, 55(11), 972–979. doi: 10.1016/j.jaac.2016.08.005

Laube, C., Lorenz, R., & van den Bos, W. (2020). Pubertal testosterone correlates with adolescent impatience and dorsal striatal activity. Developmental Cognitive Neuroscience, 42, 100749. doi: 10.1016/j.dcn.2019.100749

Lempert, K. M., Speer, M. E., Delgado, M. R., & Phelps, E. A. (2017). Positive autobiographical memory retrieval reduces temporal discounting. Social Cognitive and Affective Neuroscience, 12(10), 1584–1593. doi: 10.1093/scan/nsx086

Liu, L., Feng, T., Wang, J., & Li, H. (2012). The neural dissociation of subjective valuation from choice processes in intertemporal choice. Behavioural Brain Research, 231(1), 40–47. doi: 10.1016/j.bbr.2012.02.045

Luo, S., Ainslie, G., Giragosian, L., & Monterosso, J. R. (2009). Behavioral and Neural Evidence of Incentive Bias for Immediate Rewards Relative to Preference-Matched Delayed

Rewards. Journal of Neuroscience, 29(47), 14820–14827. doi: 10.1523/JNEUROSCI.4261-09.2009

Luo, S., Ainslie, G., Pollini, D., Giragosian, L., & Monterosso, J. R. (2012). Moderators of the association between brain activation and farsighted choice. NeuroImage, 59(2), 1469–1477. doi: 10.1016/j.neuroimage.2011.08.004

Massar, S. A. A., Libedinsky, C., Weiyan, C., Huettel, S. A., & Chee, M. W. L. (2015). Separate and overlapping brain areas encode subjective value during delay and effort discounting. NeuroImage, 120, 104–113. doi: 10.1016/j.neuroimage.2015.06.080

Mavrogiorgou, P., Enzi, B., Klimm, A.-K., Köhler, E., Roser, P., Norra, C., & Juckel, G. (2017). Serotonergic modulation of orbitofrontal activity and its relevance for decision making and impulsivity. Human Brain Mapping, 38(3), 1507–1517. doi: 10.1002/hbm.23468

McClure, S. M., Laibson, D. I., Loewenstein, G., & Cohen, J. D. (2004). Separate Neural Systems Value Immediate and Delayed Monetary Rewards. Science, 306(5695), 503–507. doi: 10.1126/science.1100907

Miedl, S. F., Wiswede, D., Marco-Pallarés, J., Ye, Z., Fehr, T., Herrmann, M., & Münte, T. F. (2015). The neural basis of impulsive discounting in pathological gamblers. Brain Imaging and Behavior, 9(4), 887–898. doi: 10.1007/s11682-015-9352-1

Murawski, C., Harris, P. G., Bode, S., D, J. F. D., & Egan, G. F. (2012). Led into Temptation? Rewarding Brand Logos Bias the Neural Encoding of Incidental Economic Decisions. PLOS ONE, 7(3), e34155. doi: 10.1371/journal.pone.0034155

Norman, L. J., Carlisi, C. O., Christakou, A., Chantiluke, K., Murphy, C., Simmons, A., … Rubia, K. (2017). Neural dysfunction during temporal discounting in paediatric Attention-Deficit/Hyperactivity Disorder and Obsessive-Compulsive Disorder. Psychiatry Research: Neuroimaging, 269, 97–105. doi: 10.1016/j.pscychresns.2017.09.008

O'Connell, G., Hsu, C.-T., Christakou, A., & Chakrabarti, B. (2018). Thinking about others and the future: Neural correlates of perspective taking relate to preferences for delayed rewards. Cognitive, Affective, & Behavioral Neuroscience, 18(1), 35–42. doi: 10.3758/s13415-017-0550-8

Peters, J., & Büchel, C. (2010). Episodic Future Thinking Reduces Reward Delay Discounting through an Enhancement of Prefrontal-Mediotemporal Interactions. Neuron, 66(1), 138–148. doi: 10.1016/j.neuron.2010.03.026

Pine, A., Seymour, B., Roiser, J. P., Bossaerts, P., Friston, K. J., Curran, H. V., & Dolan, R. J. (2009). Encoding of Marginal Utility across Time in the Human Brain. Journal of Neuroscience, 29(30), 9575–9581. doi: 10.1523/JNEUROSCI.1126-09.2009

Prévost, C., Pessiglione, M., Météreau, E., Cléry-Melin, M.-L., & Dreher, J.-C. (2010). Separate Valuation Subsystems for Delay and Effort Decision Costs. Journal of Neuroscience, 30(42), 14080–14090. doi: 10.1523/JNEUROSCI.2752-10.2010

Ripke, S., Hübner, T., Mennigen, E., Müller, K. U., Rodehacke, S., Schmidt, D., … Smolka, M. N. (2012). Reward processing and intertemporal decision making in adults and adolescents: The role of impulsivity and decision consistency. Brain Research, 1478, 36–47. doi: 10.1016/j.brainres.2012.08.034

Samanez-Larkin, G., Mata, R., Radu, P., Ballard, I., Carstensen, L., & McClure, S. (2011). Age Differences in Striatal Delay Sensitivity during Intertemporal Choice in Healthy Adults. Frontiers in Neuroscience, 5. Retrieved from https://www.frontiersin.org/articles/10.3389/fnins.2011.00126

Sasse, L. K., Peters, J., & Brassen, S. (2017). Cognitive Control Modulates Effects of Episodic Simulation on Delay Discounting in Aging. Frontiers in Aging Neuroscience, 9. Retrieved from https://www.frontiersin.org/articles/10.3389/fnagi.2017.00058

Seaman, K. L., Brooks, N., Karrer, T. M., Castrellon, J. J., Perkins, S. F., Dang, L. C., … Samanez-Larkin, G. R. (2018). Subjective value representations during effort, probability and time discounting across adulthood. Social Cognitive and Affective Neuroscience, 13(5), 449–459. doi: 10.1093/scan/nsy021

Sripada, C. S., Gonzalez, R., Phan, K. L., & Liberzon, I. (2011). The neural correlates of intertemporal decision-making: Contributions of subjective value, stimulus type, and trait impulsivity. Human Brain Mapping, 32(10), 1637–1648. doi: https://doi.org/10.1002/hbm.21136

Waegeman, A., Declerck, C. H., Boone, C., Van Hecke, W., & Parizel, P. M. (2014). Individual differences in self-control in a time discounting task: An fMRI study. Journal of Neuroscience, Psychology, and Economics, 7(2), 65–79. doi: 10.1037/npe0000018

Wang, Y., Hu, Y., Xu, J., Zhou, H., Lin, X., Du, X., & Dong, G. (2017). Dysfunctional Prefrontal Function Is Associated with Impulsivity in People with Internet Gaming Disorder during a Delay Discounting Task. Frontiers in Psychiatry, 8. Retrieved from https://www.frontiersin.org/articles/10.3389/fpsyt.2017.00287

Wiehler, A., Petzschner, F. H., Stephan, K. E., & Peters, J. (2017). Episodic Tags Enhance Striatal Valuation Signals during Temporal Discounting in pathological Gamblers. ENeuro, 4(3). doi: 10.1523/ENEURO.0159-17.2017

Wittmann, M., Leland, D. S., & Paulus, M. P. (2007). Time and decision making: Differential contribution of the posterior insular cortex and the striatum during a delay discounting task. Experimental Brain Research, 179(4), 643–653. doi: 10.1007/s00221-006-0822-y

Wittmann, M., Lovero, K. L., Lane, S. D., & Paulus, M. P. (2010). Now or later? Striatum and insula activation to immediate versus delayed rewards. Journal of Neuroscience, Psychology, and Economics, 3(1), 15–26. doi: 10.1037/a0017252

Xu, L., Liang, Z.-Y., Wang, K., Li, S., & Jiang, T. (2009). Neural mechanism of intertemporal choice: From discounting future gains to future losses. Brain Research, 1261, 65–74. doi: 10.1016/j.brainres.2008.12.061

Zhuang, J.-Y., Wang, J.-X., Lei, Q., Zhang, W., & Fan, M. (2020). Neural Basis of Increased Cognitive Control of Impulsivity During the Mid-Luteal Phase Relative to the Late Follicular Phase of the Menstrual Cycle. Frontiers in Human Neuroscience, 14. Retrieved from https://www.frontiersin.org/articles/10.3389/fnhum.2020.568399

### References: Response Inhibition


Aron, A. R., & Poldrack, R. A. (2006). Cortical and subcortical contributions to Stop signal response inhibition: Role of the subthalamic nucleus. J Neurosci, 26(9), 2424–2433. doi: 10.1523/JNEUROSCI.4682-05.2006

Bennett, D. S., Mohamed, F. B., Carmody, D. P., Bendersky, M., Patel, S., Khorrami, M., … Lewis, M. (2009). Response inhibition among early adolescents prenatally exposed to tobacco: An fMRI study. Neurotoxicology and Teratology, 31(5), 283–290. doi: 10.1016/j.ntt.2009.03.003

Berkman, E. T., Kahn, L. E., & Merchant, J. S. (2014). Training-induced changes in inhibitory control network activity. J Neurosci, 34(1), 149–157. doi: 10.1523/JNEUROSCI.3564-13.2014

Bobb, D. S., Adinoff, B., Laken, S. J., McClintock, S. M., Rubia, K., Huang, H. W., … Kozel, F. A. (2012). Neural correlates of successful response inhibition in unmedicated patients with late-life depression. Am J Geriatr Psychiatry, 20(12), 1057–1069. doi: 10.1097/JGP.0b013e318235b728

Boecker, M., Drueke, B., Vorhold, V., Knops, A., Philippen, B., & Gauggel, S. (2011). When Response Inhibition is Followed by Response Reengagement: An Event-Related fMRI Study. Human Brain Mapping, 32(1), 94–106. doi: 10.1002/hbm.21001

Boehler, C. N., Appelbaum, L. G., Krebs, R. M., Hopf, J. M., & Woldorff, M. G. (2010). Pinning down response inhibition in the brain—Conjunction analyses of the Stop-signal task. Neuroimage, 52(4), 1621–1632. doi: 10.1016/j.neuroimage.2010.04.276

Cai, W., Cannistraci, C. J., Gore, J. C., & Leung, H. C. (2014). Sensorimotor-independent prefrontal activity during response inhibition. Hum Brain Mapp, 35(5), 2119–2136. doi: 10.1002/hbm.22315

Cai, W., & Leung, H. C. (2009). Cortical activity during manual response inhibition guided by color and orientation cues. Brain Res, 1261, 20–28. doi: 10.1016/j.brainres.2008.12.073

Cai, W., & Leung, H. C. (2011). Rule-guided executive control of response inhibition: Functional topography of the inferior frontal cortex. PLoS One, 6(6), e20840. doi: 10.1371/journal.pone.0020840

Cascio, C. N., Carp, J., O'Donnell, M. B., Tinney, F. J., Jr., Bingham, C. R., Shope, J. T., … Falk, E. B. (2015). Buffering Social Influence: Neural Correlates of Response Inhibition Predict Driving Safety in the Presence of a Peer. Journal of Cognitive Neuroscience, 27(1), 83–95. doi: 10.1162/jocn_a_00693

Chen, C.-Y., Yen, J.-Y., Yen, C.-F., Chen, C.-S., Liu, G.-C., Liang, C.-Y., & Ko, C.-H. (2015). Aberrant brain activation of error processing among adults with attention deficit and hyperactivity disorder. The Kaohsiung Journal of Medical Sciences, 31(4), 179–187. doi: 10.1016/j.kjms.2015.01.001

Chevrier, A. D., Noseworthy, M. D., & Schachar, R. (2007). Dissociation of response inhibition and performance monitoring in the stop signal task using event-related fMRI. Hum Brain Mapp, 28(12), 1347–1358. doi: 10.1002/hbm.20355

Chikara, R. K., Chang, E. C., Lu, Y. C., Lin, D. S., Lin, C. T., & Ko, L. W. (2018). Monetary Reward and Punishment to Response Inhibition Modulate Activation and Synchronization Within the Inhibitory Brain Network. Front Hum Neurosci, 12, 27. doi: 10.3389/fnhum.2018.00027

Chikazoe, J., Jimura, K., Asari, T., Yamashita, K., Morimoto, H., Hirose, S., … Konishi, S. (2009). Functional dissociation in right inferior frontal cortex during performance of go/no-go task. Cereb Cortex, 19(1), 146–152. doi: 10.1093/cercor/bhn065

Chikazoe, J., Jimura, K., Hirose, S., Yamashita, K., Miyashita, Y., & Konishi, S. (2009). Preparation to inhibit a response complements response inhibition during performance of a stop-signal task. J Neurosci, 29(50), 15870–15877. doi: 10.1523/JNEUROSCI.3645-09.2009

Congdon, E., Altshuler, L. L., Mumford, J. A., Karlsgodt, K. H., Sabb, F. W., Ventura, J., … Poldrack, R. A. (2014). Neural activation during response inhibition in adult attention-deficit/hyperactivity disorder: Preliminary findings on the effects of medication and symptom severity. Psychiatry Res, 222(1–2), 17–28. doi: 10.1016/j.pscychresns.2014.02.002

Coxon, J. P., Goble, D. J., Leunissen, I., Van Impe, A., Wenderoth, N., & Swinnen, S. P. (2016). Functional Brain Activation Associated with Inhibitory Control Deficits in Older Adults. Cereb Cortex, 26(1), 12–22. doi: 10.1093/cercor/bhu165

Czapla, M., Baeuchl, C., Simon, J. J., Richter, B., Kluge, M., Friederich, H. C., … Loeber, S. (2017). Do alcohol-dependent patients show different neural activation during response inhibition than healthy controls in an alcohol-related fMRI go/no-go-task? Psychopharmacology (Berl), 234(6), 1001–1015. doi: 10.1007/s00213-017-4541-9

Dambacher, F., Sack, A. T., Lobbestael, J., Arntz, A., Brugman, S., & Schuhmann, T. (2015). Out of control: Evidence for anterior insula involvement in motor impulsivity and reactive aggression. Social Cognitive and Affective Neuroscience, 10(4), 508–516. doi: 10.1093/scan/nsu077

Durston, S., Mulder, M., Casey, B. J., Ziermans, T., & van Engeland, H. (2006). Activation in ventral prefrontal cortex is sensitive to genetic vulnerability for attention-deficit hyperactivity disorder. Biological Psychiatry, 60(10), 1062–1070. doi: 10.1016/j.biopsych.2005.12.020

Fassbender, C., Murphy, K., Foxe, J. J., Wylie, G. R., Javitt, D. C., Robertson, I. H., & Garavan, H. (2004). A topography of executive functions and their interactions revealed by functional magnetic resonance imaging. Cognitive Brain Research, 20(2), 132–143. doi: 10.1016/j.cogbrainres.2004.02.007

Fauth-Buhler, M., de Rover, M., Rubia, K., Garavan, H., Abbott, S., Clark, L., … Robbins, T. W. (2012). Brain networks subserving fixed versus performance-adjusted delay stop trials in a stop signal task. Behav Brain Res, 235(1), 89–97. doi: 10.1016/j.bbr.2012.07.023

Fedota, J. R., Hardee, J. E., Perez-Edgar, K., & Thompson, J. C. (2014). Representation of response alternatives in human presupplementary motor area: Multi-voxel pattern analysis in a go/no-go task. Neuropsychologia, 56, 110–118. doi: 10.1016/j.neuropsychologia.2013.12.022

Fuentes-Claramonte, P., Avila, C., Rodriguez-Pujadas, A., Costumero, V., Ventura-Campos, N., Bustamante, J. C., … Barros-Loscertales, A. (2016). Inferior frontal cortex activity is modulated by reward sensitivity and performance variability. Biol Psychol, 114, 127–137. doi: 10.1016/j.biopsycho.2016.01.001

Ganos, C., Kuhn, S., Kahl, U., Schunke, O., Feldheim, J., Gerloff, C., … Munchau, A. (2014). Action inhibition in Tourette syndrome. Mov Disord, 29(12), 1532–1538. doi: 10.1002/mds.25944

Garavan, H., Ross, T. J., Kaufman, J., & Stein, E. A. (2003). A midline dissociation between error-processing and response-conflict monitoring. Neuroimage, 20(2), 1132–1139. doi: 10.1016/S1053-8119(03)00334-3

Garavan, H., Ross, T. J., Murphy, K., Roche, R. A., & Stein, E. A. (2002). Dissociable executive functions in the dynamic control of behavior: Inhibition, error detection, and correction. Neuroimage, 17(4), 1820–1829. doi: 10.1006/nimg.2002.1326

Garavan, H., Ross, T. J., & Stein, E. A. (1999). Right hemispheric dominance of inhibitory control: An event-related functional MRI study. Proc Natl Acad Sci U S A, 96(14), 8301–8306. doi: 10.1073/pnas.96.14.8301

Geng, J. J., Ruff, C. C., & Driver, J. (2009). Saccades to a remembered location elicit spatially specific activation in human retinotopic visual cortex. J Cogn Neurosci, 21(2), 230–245. doi: 10.1162/jocn.2008.21025

Ghahremani, D. G., Lee, B., Robertson, C. L., Tabibnia, G., Morgan, A. T., De Shetler, N., … London, E. D. (2012). Striatal dopamine D(2)/D(3) receptors mediate response inhibition and related activity in frontostriatal neural circuitry in humans. J Neurosci, 32(21), 7316–7324. doi: 10.1523/JNEUROSCI.4284-11.2012

Halari, R., Simic, M., Pariante, C. M., Papadopoulos, A., Cleare, A., Brammer, M., … Rubia, K. (2009). Reduced activation in lateral prefrontal cortex and anterior cingulate during attention and cognitive control functions in medication-naïve adolescents with depression compared to controls. Journal of Child Psychology and Psychiatry, 50(3), 307–316. doi: 10.1111/j.1469-7610.2008.01972.x

Harle, K. M., Zhang, S., Ma, N., Yu, A. J., & Paulus, M. P. (2016). Reduced Neural Recruitment for Bayesian Adjustment of Inhibitory Control in Methamphetamine Dependence. Biol Psychiatry Cogn Neurosci Neuroimaging, 1(5), 448–459. doi: 10.1016/j.bpsc.2016.06.008

He, Q., Xiao, L., Xue, G., Wong, S., Ames, S. L., Schembre, S. M., & Bechara, A. (2014). Poor ability to resist tempting calorie rich food is linked to altered balance between neural systems involved in urge and self-control. Nutrition Journal, 13(1), 92. doi: 10.1186/1475-2891-13-92

Heitzeg, M. M., Nigg, J. T., Yau, W.-Y. W., Zucker, R. A., & Zubieta, J.-K. (2010). Striatal dysfunction marks preexisting risk and medial prefrontal dysfunction is related to problem drinking in children of alcoholics. Biological Psychiatry, 68(3), 287–295. doi: 10.1016/j.biopsych.2010.02.020

Hester, R., Foxe, J. J., Molholm, S., Shpaner, M., & Garavan, H. (2005). Neural mechanisms involved in error processing: A comparison of errors made with and without awareness. Neuroimage, 27(3), 602–608. doi: 10.1016/j.neuroimage.2005.04.035

Hester, R. L., Murphy, K., Foxe, J. J., Foxe, D. M., Javitt, D. C., & Garavan, H. (2004). Predicting success: Patterns of cortical activation and deactivation prior to response inhibition. J Cogn Neurosci, 16(5), 776–785. doi: 10.1162/089892904970726

Hester, R., Murphy, K., & Garavan, H. (2004). Beyond common resources: The cortical basis for resolving task interference. Neuroimage, 23(1), 202–212. doi: 10.1016/j.neuroimage.2004.05.024

Hough, C. M., Luks, T. L., Lai, K., Vigil, O., Guillory, S., Nongpiur, A., … Mathews, C. A. (2016). Comparison of brain activation patterns during executive function tasks in hoarding disorder and non-hoarding OCD. Psychiatry Res Neuroimaging, 255, 50–59. doi: 10.1016/j.pscychresns.2016.07.007

Hsu, J. S., Wang, P. W., Ko, C. H., Hsieh, T. J., Chen, C. Y., & Yen, J. Y. (2017). Altered brain correlates of response inhibition and error processing in females with obesity and sweet food addiction: A functional magnetic imaging study. Obes Res Clin Pract, 11(6), 677–686. doi: 10.1016/j.orcp.2017.04.011

Hughes, M. E., Fulham, W. R., Johnston, P. J., & Michie, P. T. (2012). Stop-signal response inhibition in schizophrenia: Behavioural, event-related potential and functional neuroimaging data. Biol Psychol, 89(1), 220–231. doi: 10.1016/j.biopsycho.2011.10.013

Hughes, M. E., Johnston, P. J., Fulham, W. R., Budd, T. W., & Michie, P. T. (2013). Stop-signal task difficulty and the right inferior frontal gyrus. Behav Brain Res, 256, 205–213. doi: 10.1016/j.bbr.2013.08.026

Jahfari, S., Verbruggen, F., Frank, M. J., Waldorp, L. J., Colzato, L., Ridderinkhof, K. R., & Forstmann, B. U. (2012). How preparation changes the need for top-down control of the basal ganglia when inhibiting premature actions. J Neurosci, 32(32), 10870–10878. doi: 10.1523/JNEUROSCI.0902-12.2012

Jahfari, S., Waldorp, L., Ridderinkhof, K. R., & Scholte, H. S. (2015). Visual information shapes the dynamics of corticobasal ganglia pathways during response selection and inhibition. J Cogn Neurosci, 27(7), 1344–1359. doi: 10.1162/jocn_a_00792

Jahfari, S., Waldorp, L., van den Wildenberg, W. P., Scholte, H. S., Ridderinkhof, K. R., & Forstmann, B. U. (2011). Effective connectivity reveals important roles for both the hyperdirect (fronto-subthalamic) and the indirect (fronto-striatal-pallidal) fronto-basal ganglia pathways during response inhibition. J Neurosci, 31(18), 6891–6899. doi: 10.1523/JNEUROSCI.5253-10.2011

Kaladjian, A., Jeanningros, R., Azorin, J. M., Grimault, S., Anton, J. L., & Mazzola-Pomietto, P. (2007). Blunted activation in right ventrolateral prefrontal cortex during motor response inhibition in schizophrenia. Schizophrenia Research, 97(1–3), 184–193. doi: 10.1016/j.schres.2007.07.033

Kaladjian, A., Jeanningros, R., Azorin, J. M., Nazarian, B., Roth, M., Anton, J. L., & Mazzola-Pomietto, P. (2009). Remission from mania is associated with a decrease in amygdala activation during motor response inhibition. Bipolar Disord, 11(5), 530–538. doi: 10.1111/j.1399-5618.2009.00722.x

Kaladjian, A., Jeanningros, R., Azorin, J. M., Nazarian, B., Roth, M., & Mazzola-Pomietto, P. (2009). Reduced brain activation in euthymic bipolar patients during response inhibition: An event-related fMRI study. Psychiatry Res, 173(1), 45–51. doi: 10.1016/j.pscychresns.2008.08.003

Kelly, A. M., Hester, R., Murphy, K., Javitt, D. C., Foxe, J. J., & Garavan, H. (2004). Prefrontal-subcortical dissociations underlying inhibitory control revealed by event-related fMRI. Eur J Neurosci, 19(11), 3105–3112. doi: 10.1111/j.0953-816X.2004.03429.x

Kenner, N. M., Mumford, J. A., Hommer, R. E., Skup, M., Leibenluft, E., & Poldrack, R. A. (2010). Inhibitory motor control in response stopping and response switching. J Neurosci, 30(25), 8512–8518. doi: 10.1523/JNEUROSCI.1096-10.2010

Kiehl, K. A., Liddle, P. F., & Hopfinger, J. B. (2000). Error processing and the rostral anterior cingulate: An event-related fMRI study. Psychophysiology, 37(2), 216–223.

Ko, C. H., Hsieh, T. J., Chen, C. Y., Yen, C. F., Chen, C. S., Yen, J. Y., … Liu, G. C. (2014). Altered brain activation during response inhibition and error processing in subjects with Internet gaming disorder: A functional magnetic imaging study. Eur Arch Psychiatry Clin Neurosci, 264(8), 661–672. doi: 10.1007/s00406-013-0483-3

Ko, L.-W. ; S. (2016). Neural mechanisms of inhibitory response in a battlefield scenario: A silmutaneous fMRI-EEG study. Frontiers in Human Neuroscience, 10, 1–15.

Kohler, S., Schumann, A., de la Cruz, F., Wagner, G., & Bar, K. J. (2018). Towards response success prediction: An integrative approach using high-resolution fMRI and autonomic indices. Neuropsychologia, 119, 182–190. doi: 10.1016/j.neuropsychologia.2018.08.003

Lavallee, C. F., Herrmann, C. S., Weerda, R., & Huster, R. J. (2014). Stimulus-response mappings shape inhibition processes: A combined EEG-fMRI study of contextual stopping. PLoS One, 9(4), e96159. doi: 10.1371/journal.pone.0096159

Lee, N. C., Weeda, W. D., Insel, C., Somerville, L. H., Krabbendam, L., & Huizinga, M. (2018). Neural substrates of the influence of emotional cues on cognitive control in risk-taking adolescents. Developmental Cognitive Neuroscience, 31, 20–34. doi: 10.1016/j.dcn.2018.04.007

Liddle, P. F., Kiehl, K. A., & Smith, A. M. (2001). Event-related fMRI study of response inhibition. Hum Brain Mapp, 12(2), 100–109.

Lim, L., Hart, H., Mehta, M. A., Simmons, A., Mirza, K., & Rubia, K. (2015). Neural Correlates of Error Processing in Young People With a History of Severe Childhood Abuse: An fMRI Study. American Journal of Psychiatry, 172(9), 892–900. doi: 10.1176/appi.ajp.2015.14081042

Lock, J., Garrett, A., Beenhakker, J., & Reiss, A. L. (2011). Aberrant brain activation during a response inhibition task in adolescent eating disorder subtypes. The American Journal of Psychiatry, 168(1), 55–64. doi: 10.1176/appi.ajp.2010.10010056

Lorenz, R. C., Gleich, T., Buchert, R., Schlagenhauf, F., Kuhn, S., & Gallinat, J. (2015). Interactions between glutamate, dopamine, and the neuronal signature of response inhibition in the human striatum. Hum Brain Mapp, 36(10), 4031–4040. doi: 10.1002/hbm.22895

Marco-Pallares, J., Camara, E., Munte, T. F., & Rodriguez-Fornells, A. (2008). Neural mechanisms underlying adaptive actions after slips. J Cogn Neurosci, 20(9), 1595–1610. doi: 10.1162/jocn.2008.20117

Mazzola-Pomietto, P., Kaladjian, A., Azorin, J. M., Anton, J. L., & Jeanningros, R. (2009). Bilateral decrease in ventrolateral prefrontal cortex activation during motor response inhibition in mania. J Psychiatr Res, 43(4), 432–441. doi: 10.1016/j.jpsychires.2008.05.004

Mohammadi, B., Kollewe, K., Cole, D. M., Fellbrich, A., Heldmann, M., Samii, A., … Kramer, U. M. (2015). Amyotrophic lateral sclerosis affects cortical and subcortical activity underlying motor inhibition and action monitoring. Hum Brain Mapp, 36(8), 2878–2889. doi: 10.1002/hbm.22814

Montojo, C. A., Jalbrzikowski, M., Congdon, E., Domicoli, S., Chow, C., Dawson, C., … Bearden, C. E. (2015). Neural substrates of inhibitory control deficits in 22q11.2 deletion syndrome. Cereb Cortex, 25(4), 1069–1079. doi: 10.1093/cercor/bht304

Mulder, M. J., Baeyens, D., Davidson, M. C., Casey, B. J., DEN Ban, E. V., VAN Engeland, H., & Durston, S. (2008). Familial vulnerability to ADHD affects activity in the cerebellum in addition to the prefrontal systems. Journal of the American Academy of Child and Adolescent Psychiatry, 47(1), 68–75. doi: 10.1097/chi.0b013e31815a56dc

Nakata, H., Sakamoto, K., Ferretti, A., Gianni Perrucci, M., Del Gratta, C., Kakigi, R., & Romani, G. L. (2008). Executive functions with different motor outputs in somatosensory Go/Nogo tasks: An event-related functional MRI study. Brain Res Bull, 77(4), 197–205. doi: 10.1016/j.brainresbull.2008.07.008

Nosarti, C., Rubia, K., Smith, A. B., Frearson, S., Williams, S. C., Rifkin, L., & Murray, R. M. (2006). Altered functional neuroanatomy of response inhibition in adolescent males who were born very preterm. Developmental Medicine and Child Neurology, 48(4), 265–271. doi: 10.1017/S0012162206000582

O'Connor, D. A., Rossiter, S., Yucel, M., Lubman, D. I., & Hester, R. (2012). Successful inhibitory control over an immediate reward is associated with attentional disengagement in visual processing areas. Neuroimage, 62(3), 1841–1847. doi: 10.1016/j.neuroimage.2012.05.040

Rae, C. L., Hughes, L. E., Weaver, C., Anderson, M. C., & Rowe, J. B. (2014). Selection and stopping in voluntary action: A meta-analysis and combined fMRI study. Neuroimage, 86, 381–391. doi: 10.1016/j.neuroimage.2013.10.012

Rodriguez-Pujadas, A., Sanjuan, A., Fuentes, P., Ventura-Campos, N., Barros-Loscertales, A., & Avila, C. (2014). Differential neural control in early bilinguals and monolinguals during response inhibition. Brain Lang, 132, 43–51. doi: 10.1016/j.bandl.2014.03.003

Rothmayr, C., Sodian, B., Hajak, G., Dohnel, K., Meinhardt, J., & Sommer, M. (2011). Common and distinct neural networks for false-belief reasoning and inhibitory control. Neuroimage, 56(3), 1705–1713. doi: 10.1016/j.neuroimage.2010.12.052

Rubia, K., Smith, A. B., Brammer, M. J., & Taylor, E. (2003). Right inferior prefrontal cortex mediates response inhibition while mesial prefrontal cortex is responsible for error detection. Neuroimage, 20(1), 351–358. doi: 10.1016/s1053-8119(03)00275-1

Rubia, Katya, Lim, L., Ecker, C., Halari, R., Giampietro, V., Simmons, A., … Smith, A. (2013). Effects of age and gender on neural networks of motor response inhibition: From adolescence to mid-adulthood. NeuroImage, 83, 690–703. doi: 10.1016/j.neuroimage.2013.06.078

Rubia, Katya, Smith, A. B., Woolley, J., Nosarti, C., Heyman, I., Taylor, E., & Brammer, M. (2006). Progressive increase of frontostriatal brain activation from childhood to adulthood during event-related tasks of cognitive control. Human Brain Mapping, 27(12), 973–993. doi: 10.1002/hbm.20237

Schel, M. A., Kuhn, S., Brass, M., Haggard, P., Ridderinkhof, K. R., & Crone, E. A. (2014). Neural correlates of intentional and stimulus-driven inhibition: A comparison. Front Hum Neurosci, 8, 27. doi: 10.3389/fnhum.2014.00027

Schulz, K. P., Fan, J., Tang, C. Y., Newcorn, J. H., Buchsbaum, M. S., Cheung, A. M., & Halperin, J. M. (2004). Response Inhibition in Adolescents Diagnosed With Attention Deficit Hyperactivity Disorder During Childhood: An Event-Related fMRI Study. American Journal of Psychiatry, 161(9), 1650–1657. doi: 10.1176/appi.ajp.161.9.1650

Sebastian, A., Baldermann, C., Feige, B., Katzev, M., Scheller, E., Hellwig, B., … Kloppel, S. (2013). Differential effects of age on subcomponents of response inhibition. Neurobiol Aging, 34(9), 2183–2193. doi: 10.1016/j.neurobiolaging.2013.03.013

Sebastian, A., Gerdes, B., Feige, B., Kloppel, S., Lange, T., Philipsen, A., … Tuscher, O. (2012). Neural correlates of interference inhibition, action withholding and action cancelation in adult ADHD. Psychiatry Res, 202(2), 132–141. doi: 10.1016/j.pscychresns.2012.02.010

Sebastian, A., Pohl, M. F., Kloppel, S., Feige, B., Lange, T., Stahl, C., … Tuscher, O. (2013). Disentangling common and specific neural subprocesses of response inhibition. Neuroimage, 64, 601–615. doi: 10.1016/j.neuroimage.2012.09.020

Sebastian, A., Rossler, K., Wibral, M., Mobascher, A., Lieb, K., Jung, P., & Tuscher, O. (2017). Neural Architecture of Selective Stopping Strategies: Distinct Brain Activity Patterns Are Associated with Attentional Capture But Not with Outright Stopping. J Neurosci, 37(40), 9785–9794. doi: 10.1523/JNEUROSCI.1476-17.2017

Sharp, D. J., Bonnelle, V., De Boissezon, X., Beckmann, C. F., James, S. G., Patel, M. C., & Mehta, M. A. (2010). Distinct frontal systems for response inhibition, attentional capture, and error processing. Proc Natl Acad Sci U S A, 107(13), 6106–6111. doi: 10.1073/pnas.1000175107

Steele, V. R., Claus, E. D., Aharoni, E., Harenski, C., Calhoun, V. D., Pearlson, G., & Kiehl, K. A. (2014). A large scale (N=102) functional neuroimaging study of error processing in a Go/NoGo task. Behav Brain Res, 268, 127–138. doi: 10.1016/j.bbr.2014.04.001

Swann, N. C., Cai, W., Conner, C. R., Pieters, T. A., Claffey, M. P., George, J. S., … Tandon, N. (2012). Roles for the pre-supplementary motor area and the right inferior frontal gyrus in stopping action: Electrophysiological responses and functional and structural connectivity. Neuroimage, 59(3), 2860–2870. doi: 10.1016/j.neuroimage.2011.09.049

Tabu, H., Mima, T., Aso, T., Takahashi, R., & Fukuyama, H. (2011). Functional relevance of pre-supplementary motor areas for the choice to stop during Stop signal task. Neurosci Res, 70(3), 277–284. doi: 10.1016/j.neures.2011.03.007

Tabu, H., Mima, T., Aso, T., Takahashi, R., & Fukuyama, H. (2012). Common inhibitory prefrontal activation during inhibition of hand and foot responses. Neuroimage, 59(4), 3373–3378. doi: 10.1016/j.neuroimage.2011.10.092

van der Meer, L., Groenewold, N. A., Pijnenborg, M., & Aleman, A. (2013). Psychosis-proneness and neural correlates of self-inhibition in theory of mind. PLoS One, 8(7), e67774. doi: 10.1371/journal.pone.0067774

van Eijk, J., Sebastian, A., Krause-Utz, A., Cackowski, S., Demirakca, T., Biedermann, S. V., … Tuscher, O. (2015). Women with borderline personality disorder do not show altered BOLD responses during response inhibition. Psychiatry Res, 234(3), 378–389. doi: 10.1016/j.pscychresns.2015.09.017

Walther, S., Goya-Maldonado, R., Stippich, C., Weisbrod, M., & Kaiser, S. (2010). A supramodal network for response inhibition. Neuroreport, 21(3), 191–195. doi: 10.1097/WNR.0b013e328335640f

White, T. P., Loth, E., Rubia, K., Krabbendam, L., Whelan, R., Banaschewski, T., … IMAGEN Consortium. (2014). Sex differences in COMT polymorphism effects on prefrontal inhibitory control in adolescence. Neuropsychopharmacology: Official Publication of the American College of Neuropsychopharmacology, 39(11), 2560–2569. doi: 10.1038/npp.2014.107

Wilbertz, T., Deserno, L., Horstmann, A., Neumann, J., Villringer, A., Heinze, H. J., … Schlagenhauf, F. (2014). Response inhibition and its relation to multidimensional impulsivity. Neuroimage, 103, 241–248. doi: 10.1016/j.neuroimage.2014.09.021

Xu, B., Levy, S., Butman, J., Pham, D., Cohen, L. G., & Sandrini, M. (2015). Effect of foreknowledge on neural activity of primary ‘go’ responses relates to response stopping and switching. Front Hum Neurosci, 9, 34. doi: 10.3389/fnhum.2015.00034

Xu, K. Z., Anderson, B. A., Emeric, E. E., Sali, A. W., Stuphorn, V., Yantis, S., & Courtney, S. M. (2017). Neural Basis of Cognitive Control over Movement Inhibition: Human fMRI and

Primate Electrophysiology Evidence. Neuron, 96(6), 1447-1458 e6. doi: 10.1016/j.neuron.2017.11.010

Xue, G., Aron, A. R., & Poldrack, R. A. (2008). Common neural substrates for inhibition of spoken and manual responses. Cereb Cortex, 18(8), 1923–1932. doi: 10.1093/cercor/bhm220

Zandbelt, B. B., van Buuren, M., Kahn, R. S., & Vink, M. (2011). Reduced proactive inhibition in schizophrenia is related to corticostriatal dysfunction and poor working memory. Biol Psychiatry, 70(12), 1151–1158. doi: 10.1016/j.biopsych.2011.07.028

Zandbelt, B. B., & Vink, M. (2010). On the role of the striatum in response inhibition. PLoS One, 5(11), e13848. doi: 10.1371/journal.pone.0013848

Zheng, D., Oka, T., Bokura, H., & Yamaguchi, S. (2008). The key locus of common response inhibition network for no-go and stop signals. J Cogn Neurosci, 20(8), 1434–1442. doi: 10.1162/jocn.2008.20100
