## Supplementary methods for "Charting the brain networks of impulsivity: Meta-analytic synthesis, functional connectivity modelling and neurotransmitter associations"

### Supplementary Methods to Gell et al., 2022

#### 1. Main Effect Thresholding and Masking

In order to account for the higher power of the controlled action > baseline/go meta-analysis (i.e. large discrepancy in the number of included studies: 26 in impulsive action vs 103 in controlled action) we used a subsampling procedure. This was achieved by iteratively sub-sampling 26 experiments from the full controlled action database 10 000 times. In each iteration a meta-analysis was calculated and for each voxel an absence or presence of a significant main effect was recorded. Thus a frequency of significant results across the 10 000 iterations for each voxel was obtained. The resulting probabilistic map represents the number of times each voxel participated in a main effect over the permutation procedure as a proportion of the total number of permutations. Finally, this probabilistic map was thresholded at the 90th percentile to remove voxels with a very low probability of participating in a main effect and used to mask the controlled choice main effect map before computing the contrast (see figure x in supplement for illustration of this procedure). An illustration of this procedure is displayed below.

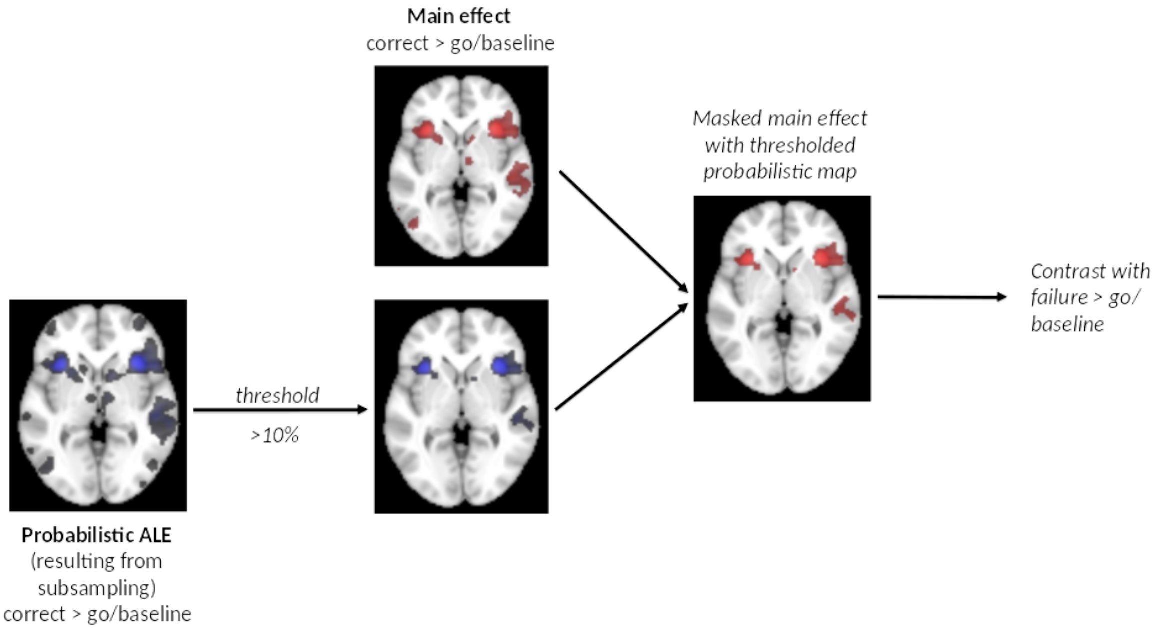


Figure S1. Main effect masking procedure

#### 2. fMRIPrep preprocessing

##### Anatomical data preprocessing

T1-weighted (T1w) images were corrected for intensity non-uniformity (INU) with `N4BiasFieldCorrection` (Tustison et al., 2010), distributed with ANTs 2.2.0 (Avants, Epstein,

Grossman, and Gee, 2008). The T1w-reference was then skull-stripped with a *Nipype* implementation of the `antsBrainExtraction.sh` workflow (from ANTs), using OASIS30ANTs as target template. Brain tissue segmentation of cerebrospinal fluid (CSF), white-matter (WM) and gray-matter (GM) was performed on the brain-extracted T1w using `fast` [FSL 5.0.9, Zhang, Brady, and Smith, 2001)].

A T1w-reference map was computed after registration of 2 T1w images (after INU-correction) using `mri_robust_template` [FreeSurfer 6.0.1, Reuter, Rosas, and Fischl, 2010)]. Brain surfaces were reconstructed using `recon-all` [FreeSurfer 6.0.1, Dale, Fischl, and Sereno, 1999)], and the brain mask estimated previously was refined with a custom variation of the method to reconcile ANTs-derived and FreeSurfer-derived segmentations of the cortical gray-matter of Mindboggle (Klein et al., 2017). Volume-based spatial normalization to two standard spaces (MNI152NLin6Asym, MNI152NLin2009cAsym) was performed through nonlinear registration with `antsRegistration` (ANTs 2.2.0), using brain-extracted versions of both T1w reference and the T1w template. The following templates were selected for spatial normalization: *FSL's MNI ICBM 152 non-linear 6th Generation Asymmetric Average Brain Stereotaxic Registration Model* [Evans, Janke, Collins, and Baillet, 2012), RRID:SCR_002823; TemplateFlow ID: MNI152NLin6Asym], *ICBM 152 Nonlinear Asymmetrical template version 2009c* [Fonov, Evans, McKinstry, Almli and Collins, et al., 2009), RRID:SCR_008796; TemplateFlow ID: MNI152NLin2009cAsym].

#### Functional data preprocessing

For the BOLD runs found per subject (across all tasks and sessions), the following preprocessing was performed. First, a reference volume and its skull-stripped version were generated using a custom methodology of *fMRIPrep*. Head-motion parameters with respect to the BOLD reference (transformation matrices, and six corresponding rotation and translation parameters) are estimated before any spatiotemporal filtering using `mcflirt` [FSL 5.0.9, (Jenkinson, Bannister, Brady, and Smith, 2002)]. BOLD runs were slice-time corrected using `3dTshift` from AFNI 20160207 (Cox and Hyde, 1997). Susceptibility distortion correction (SDC) was omitted. The BOLD reference was then co-registered to the T1w reference using `bbregister` (FreeSurfer) which implements boundary-based registration (Greve and Fischl, 2009). Co-registration was configured with six degrees of freedom. The BOLD time-series were resampled onto the following surfaces (FreeSurfer reconstruction nomenclature): *fsaverage*.

The BOLD time-series (including slice-timing correction when applied) were resampled onto their original, native space by applying the transforms to correct for head-motion. These resampled BOLD time-series will be referred to as *preprocessed BOLD in original space*, or just *preprocessed BOLD*. The BOLD time-series were resampled into standard space, generating a *preprocessed BOLD run in MNI152NLin6Asym space*. First, a reference volume and its skull-stripped version were generated using a custom methodology of *fMRIPrep*. *Grayordinates* files (Glasser et al., 2013) containing 91k samples were also generated using the highest-resolution ``fsaverage`` as intermediate standardized surface space.

Automatic removal of motion artifacts using independent component analysis [ICA-AROMA, Pruim et al., (2015)] was performed on the *preprocessed BOLD on MNI space* time-series after removal of non-steady state volumes and spatial smoothing with an isotropic, Gaussian kernel of 6mm FWHM (full-width half-maximum). Corresponding "non-aggresively" denoised runs were produced after such Smoothing. Additionally, the "aggressive" noise-regressors were collected and placed in the corresponding confounds file. Several confounding time-series were calculated based on the *preprocessed BOLD*: framewise displacement (FD), DVARS and three region-wise global signals. FD was computed using two formulations following Power (absolute sum of relative motions, Power et al., (2014)) and Jenkinson (relative root mean square displacement between affines, Jenkinson et al. (2002)). FD and DVARS are calculated for each functional run, both using their implementations in *Nipype* [following the definitions by Power et al., (2014)]. The three global signals are extracted within the CSF, the WM, and the whole-brain masks.
Additionally, a set of physiological regressors were extracted to allow for component-based noise correction [*CompCor*, (Behzadi, Restom, Liau and Liu, 2007)]. Principal components are estimated after high-pass filtering the *preprocessed BOLD* time-series (using a discrete cosine filter with 128s cut-off) for the two *CompCor* variants: temporal (tCompCor) and anatomical (aCompCor). tCompCor components are then calculated from the top 5% variable voxels within a mask covering the subcortical regions. This subcortical mask is obtained by heavily eroding the brain mask, which ensures it does not include cortical GM regions. For aCompCor, components are calculated within the intersection of the aforementioned mask and the union of CSF and WM masks calculated in T1w space, after their projection to the native space of each functional run (using the inverse BOLD-to-T1w transformation). Components are also calculated separately within the WM and CSF masks. For each CompCor decomposition, the *k* components with the largest singular values are retained, such that the retained components' time series are sufficient to explain 50 percent of variance across the nuisance mask (CSF, WM, combined, or temporal). The remaining components are dropped from consideration.

The head-motion estimates calculated in the correction step were also placed within the corresponding confounds file. The confound time series derived from head motion estimates and global signals were expanded with the inclusion of temporal derivatives and quadratic terms for each (Satterthwaite et al., 2013). Frames that exceeded a threshold of 0.5 mm FD or 1.5 standardised DVARS were annotated as motion outliers. All resamplings can be performed with *a single interpolation step* by composing all the pertinent transformations (i.e. head-motion transform matrices, susceptibility distortion correction when available, and co-registrations to anatomical and output spaces). Gridded (volumetric) resamplings were performed using `antsApplyTransforms` (ANTs), configured with Lanczos interpolation to minimize the smoothing effects of other kernels (Lanczos, 1964). Non-gridded (surface) resamplings were performed using `mri_vol2surf` (FreeSurfer).

Many internal operations of *fMRIPrep* use *Nilearn* 0.6.2 (Abraham et al., 2014), mostly within the functional processing workflow. For more details of the pipeline, see [the section corresponding to workflows in *fMRIPrep*'s documentation](https://fmriprep.readthedocs.io/en/latest/workflows.html "FMRIPrep's documentation").

Copyright Waiver

The above boilerplate text was automatically generated by fMRIPrep with the express intention that users should copy and paste this text into their manuscripts *unchanged*. It is released under the [CC0](https://creativecommons.org/publicdomain/zero/1.0/) license.
