## Supplementary results for "Charting the brain networks of impulsivity: Meta-analytic synthesis, functional connectivity modelling and neurotransmitter associations"

### Supplementary Figures and Tables to Gell et al., 2022

#### 1. Supplementary Figures

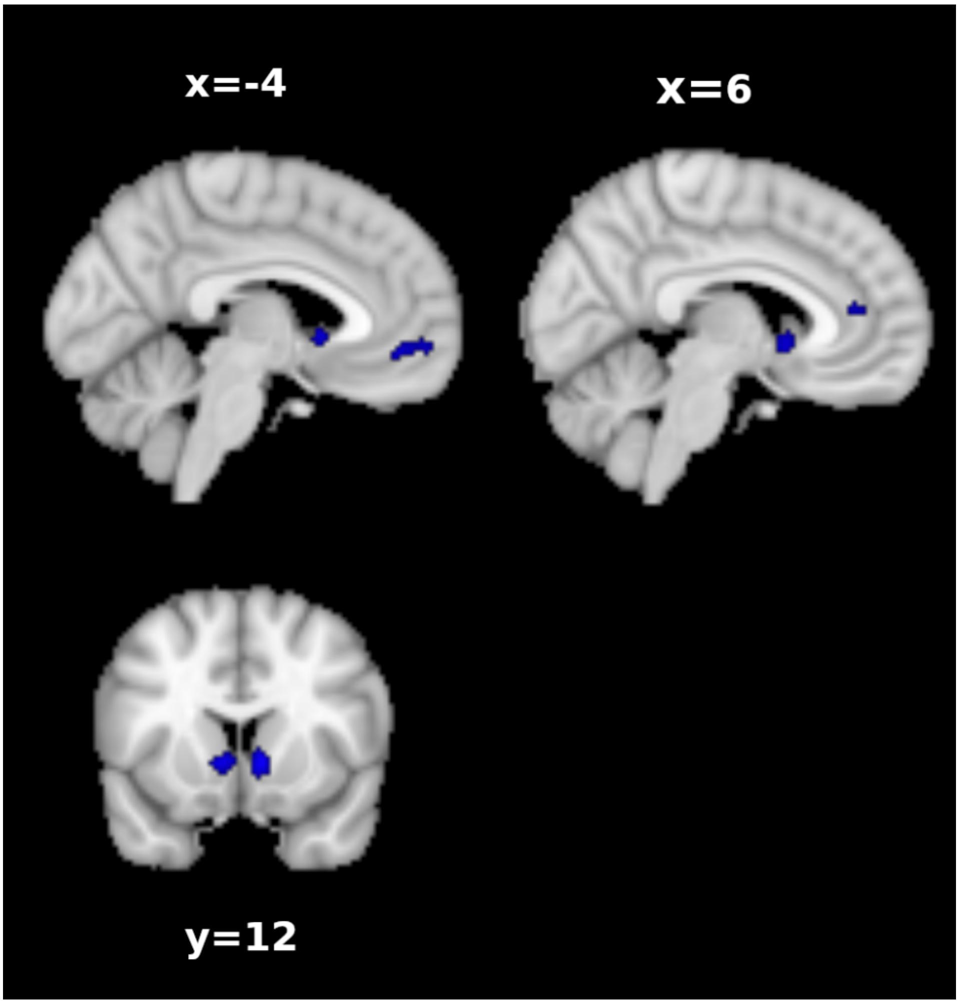


Supplementary Figure 1. Conjunction of impulsive responding and subjective value meta-analyses in DCS.

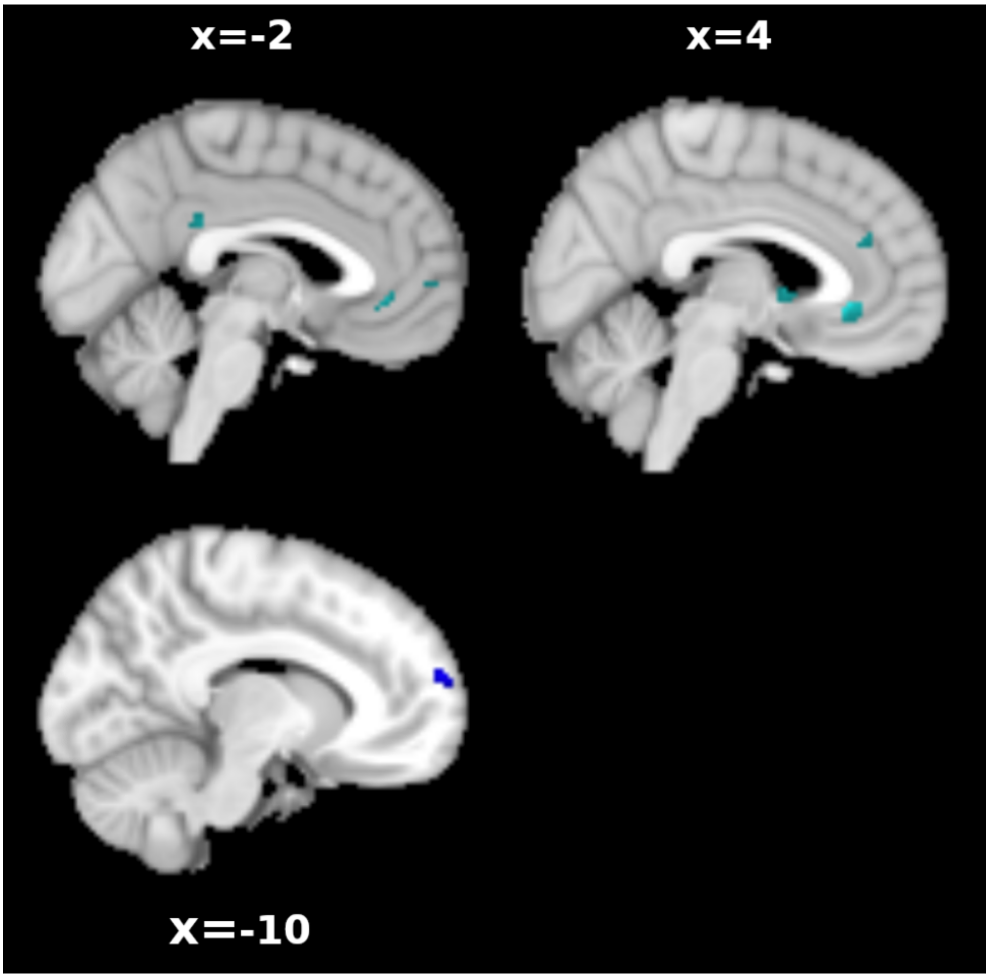


Supplementary Figure 2. Contrast analysis of impulsive responding and subjective value meta-analyses in DCS. Impulsive responding > subjective value in blue and opposite contrast in green.

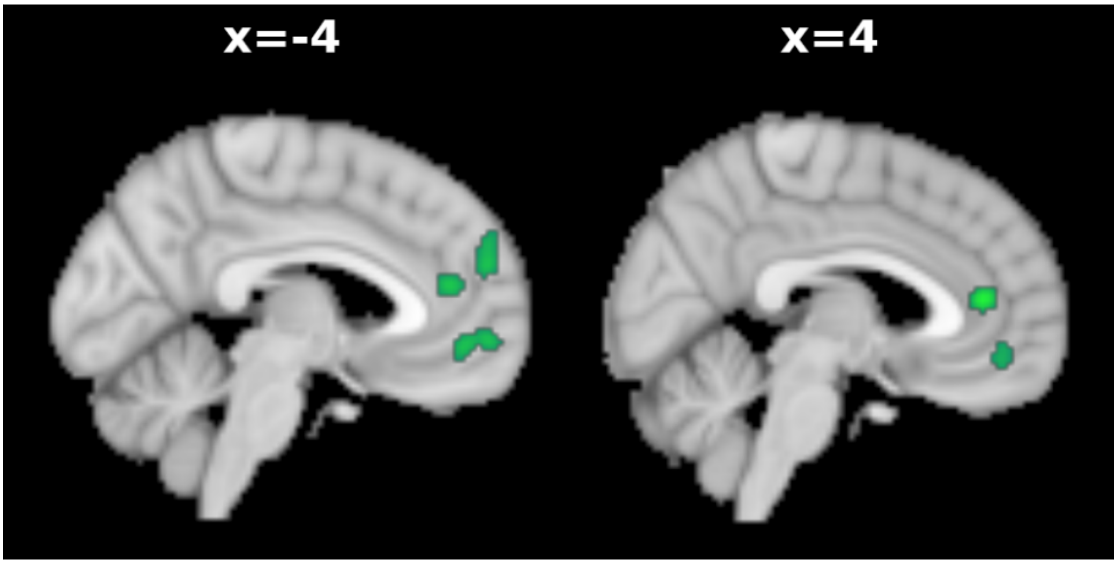


Supplementary Figure 3. Meta-analysis of impulsive responding contrast in DCS without studies correlating activity with k.

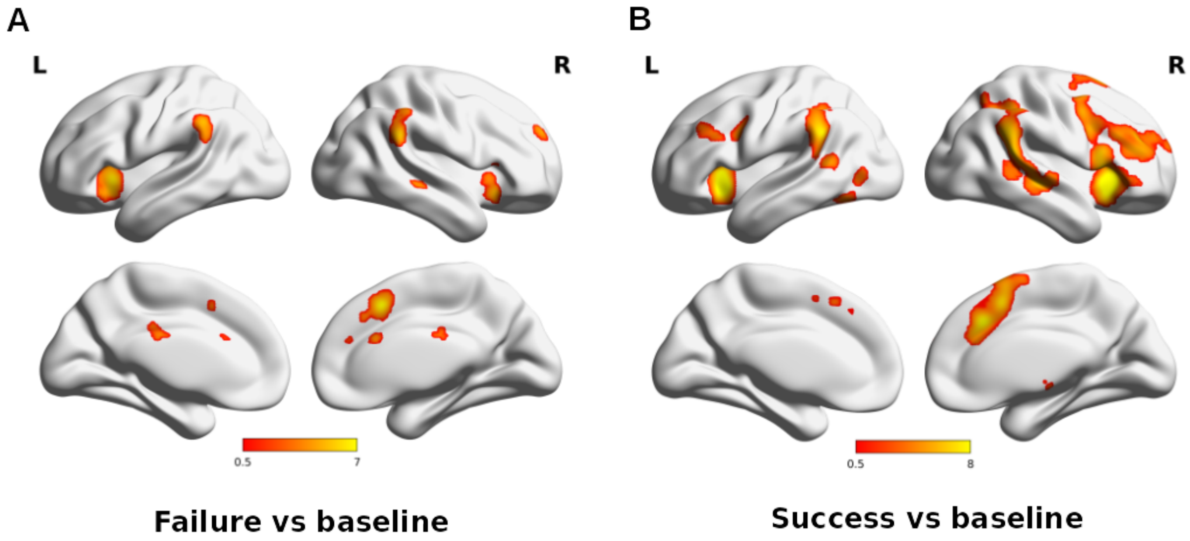


Supplementary Figure 4. Results of failure of inhibition against baseline (A) and successful inhibition against baseline (B) meta-analysis.

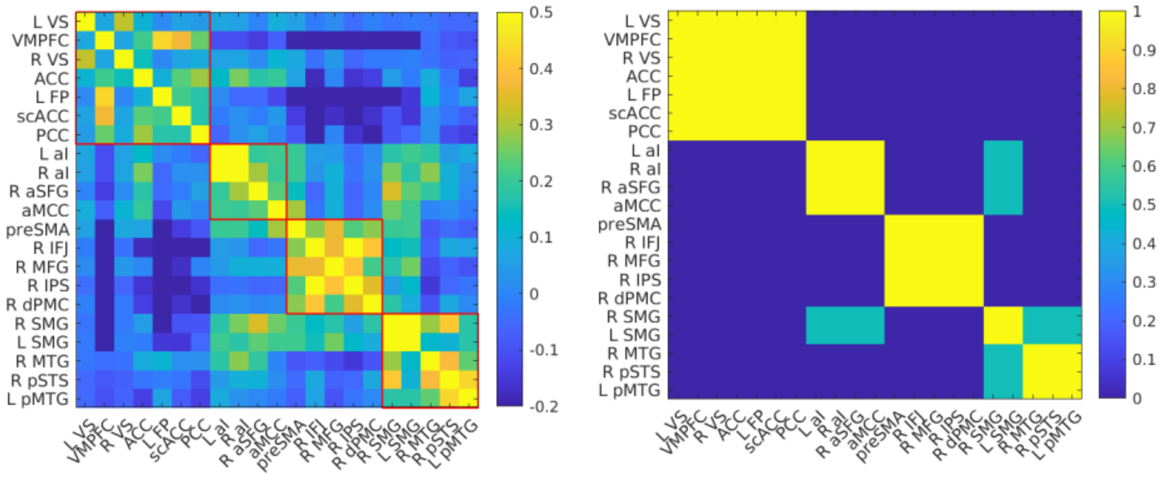


Supplementary Figure 5. Community detection results in the replication sample. Resulting communities on the left and agreement matrix on the right side (1 = all 1000 repeats yielded the same clustering solution).

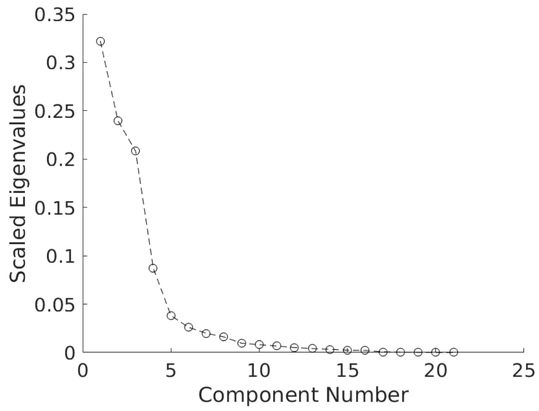


Supplementary Figure 6. Scree plot showing the eigenvalues of each component.

### Supplementary Tables

#### Supplementary table 1: Checklist for Neuroimaging Meta-Analyses by Müller et al. (2018)

| | |
|---|---|
| The research question was specifically defined | YES, and it included the following contrasts:<br><br>1) Impulsive choice (smaller sooner > larger later, immediate > delay, β > δ, correlation with k)<br>2) Controlled choice (larger later > smaller sooner, delay > immediate, δ > β, correlation with k)<br>3) Subjective value (parametric modulation of and correlation with subjective value)<br>4) Impulsive action (commission error > go/baseline/rest)<br>5) Controlled action (successful inhibition > go/baseline/rest)<br><br>The specific contrasts are reports in the method section |
| The literature search was systematic | YES, it included the following keywords in the following databases:<br><br>1. Delay consequence sensitivity<br>“delay discounting” or “temporal discounting” or “delayed reward” and “fMRI” or “functional magnetic resonance imaging”<br><br>2. Response inhibition<br>“stop signal task” or “go nogo task” or “response inhibition” or “inhibition” or "action withholding" or "action cancellation" or "action inhibition" or "motor inhibition" or "inhibitory control" and “fMRI” or “functional magnetic resonance imaging”<br><br>Databases: PubMed, Web of Science |
| Detailed inclusion and exclusion criteria were applied | YES, and reasons of non-standard criteria were:<br><br>Inclusion of: fMRI studies, healthy participants, mean age >= 12, contrast of interest, no region of interest, no pharmacological interventions or connectivity-based analyses |
| Sample overlap was taken into account | YES, using the following method:<br><br>In cases of partly overlapping subject groups (e.g. as in the case of Kable 2007 and Kable 2010), coordinates were pooled to form a single experiment and the smaller N of the two original experiments was used as the input to ALE. |

| | |
|---|---|
| All experiments used the same search coverage (state how brain coverage was assessed and how small volume corrections and conjunctions were taken into account) | YES, the search coverage was the following:<br><br>- whole-brain coverage only<br>- exclusion of ROI studies |
| Studies are converted to a common reference space | YES, using the following conversion(s):<br><br>Coordinates reported in Talairach space were converted to MNI space (Lancaster et al., 2007) |
| Data extraction was conducted by two investigators (ideal case) or double-checked by the same investigator (state how double-checking was performed) | YES, the following authors:<br><br>- MG, VM, EC checked inclusion criteria<br>- MG extracted coordinates<br>- MG and VK extracted other info: age, sex, sample size, contrast, space<br>- VK and VM double-checked the following data: inclusion criteria, extracted coordinates, all other infos |
| The paper includes a table with at least the references, basic study description (e.g., for fMRI tasks, stimuli), contrasts and basic sample descriptions (e.g., size, mean age and gender distribution, specific characteristics) of the included studies, source of information (e.g., contact with authors), reference space | YES, and also the following data:<br><br>- If further information was received by the authors |
| The study protocol and all analyses was planned before- hand, including the methods and parameters used for inference, correction for multiple testing, etc. | YES. The meta-analysis was not pre-registered; however, all analyses including methods and recommended parameters used for inference were planned before starting the literature search |
| The paper includes meta-analytic diagnostics | No diagnostics are reported given that all analyses included at least 21 experiments and a cFWE correction. As shown in Eickhoff 2016 results are robust with regard to being driven by only a few experiments. |
